# DeepTCR: a deep learning framework for understanding T-cell receptor sequence signatures within complex T-cell repertoires

**DOI:** 10.1101/464107

**Authors:** John-William Sidhom, H. Benjamin Larman, Petra Ross-MacDonald, Megan Wind-Rotolo, Drew M. Pardoll, Alexander S. Baras

## Abstract

Deep learning algorithms have been utilized to achieve enhanced performance in pattern-recognition tasks, such as in image and vocal recognition ^1,2^. The ability to learn complex patterns in data has tremendous implications in the genomics and immunology worlds, where sequence motifs become learned ‘features’ that can be used to predict functionality, guiding our understanding of disease and basic biology ^3–6^. T-cell receptor (TCR) sequencing assesses the diversity of the adaptive immune system, where complex structural patterns in the TCR can be used to model its antigenic interaction. We present DeepTCR, a broad collection of unsupervised and supervised deep learning methods able to uncover structure in highly complex and large TCR sequencing data by learning a joint representation of a given TCR by its CDR3 sequences, V/D/J gene usage, and HLA background in which the T-cells reside. We demonstrate the utility of deep learning to provide an improved ‘featurization’ of the TCR across multiple human and murine datasets, including improved classification of antigen-specific TCR’s in both unsupervised and supervised learning tasks, understanding immunotherapy-related shaping of repertoire in the murine setting, and predicting response to checkpoint blockade immunotherapy from pre-treatment tumor biopsies in a clinical trial of melanoma. Our results show the flexibility and capacity for deep neural networks to handle the complexity of high-dimensional TCR genomic data for both descriptive and predictive purposes across basic science and clinical research.

---

Next-Generation Sequencing (NGS) has allowed a comprehensive description and understanding of the complexity encoded at the genomic level in a wide variety of organisms. The applications of NGS have grown rapidly as this technology has become a molecular microscope for understanding the genomic basis for the fundamental functions of the cell^7^. In parallel to this explosion of NGS applications, in the machine learning world, deep learning has seen a similar expansion of applications as computational resources have grown; there exists many opportunities to apply deep learning in genomics as the data generated from NGS is very large and highly complex.

T-cell receptor sequencing (TCR-Seq) is an application of NGS that has allowed scientists across many disciplines to characterize the diversity of the adaptive immune response^8–18^. By selectively amplifying and sequencing the highly diverse antigen-specific CDR3 region of the β-chain of the T-cell receptor, scientists have been able to study clonal expansion as a probe for responses to both foreign and native potential antigens^19^. With this new sequencing technology, there has arisen a need to develop analytical tools to parse and draw meaningful concepts from the data, since antigen-specific T-cells exist within a sea of T-cells with specificities irrelevant to the microbe or tumor cell being assessed. In recent work, investigators have applied conventional sequence analytics, where either targeted motif searches or sequence alignment algorithms have been used to begin parsing the structural data within TCR-Seq^20–22^. However, identifying signal over noise is particularly challenging in studying *in-vivo* T-cell responses such as tumor-specific T-cell responses, which appear to be mediated by a small proportion of tumor-infiltrating lymphocytes (TIL) and peripheral blood lymphocytes (PBL)^23–25^. While these CDR3 alignment algorithms have been used successfully to assign TCRs to a limited number of antigens after multimer sorting, they have done so in absence of the 100-1000x background of irrelevant specificities seen in typical *in-vivo* T-cell responses^21,26^.

In light of this need to better ‘featurize’ TCR sequences, we turned to deep learning primarily through the use of convolutional neural networks (CNN’s) as a powerful means to extract important features from sequencing data for both descriptive and predictive purposes. As has been demonstrated in previous genomic applications of deep learning, the main advantage of CNN’s in this application is the ability to learn these sequence motifs (referred to as kernels in this context) through some objective function given to the network^3^. These learned motifs can then be used as part of a complex deep learning model to either describe the data in a new latent space or be used for a classification task.

We present DeepTCR, a platform of both unsupervised and supervised deep learning that is able to be applied at the level of individual T-cell receptor sequences as well as at the level of whole T-cell repertoires, which can learn patterns in the data that may be used for both descriptive and predictive purposes. In order to demonstrate the utility of these algorithms, we collected two types of datasets (Supplementary Fig. 1); samples sorted by antigen-specificity (*Glanville_2017, Sidhom_2017, Dash_2017, Zhang_2018, 10x_Genomics),* and samples taken from a heterogenous immune response with unknown specificities including cohorts of tumor-bearing mice treated with various immunotherapies *(Rudqvist_2017)* and HLA-typed melanoma patients whose tumor biopsies were TCRβ CDR3 sequenced prior to the initiation of checkpoint blockade immunotherapy *(CheckMate-038)*^20,22,27–29^. The antigen-specific datasets represent experiments where T-cells were tetramer-sorted for a specific antigen and where the ground truth label corresponds to a particular antigen-specificity. In contrast, the first immunotherapy dataset consists of a murine experiment where 4 syngeneic cohorts of tumor-bearing mice were treated with various therapeutic interventions (Control = No Intervention, RT = Radiation Therapy, 9H10 = α-CTLA4, Combo = Radiation Therapy + α-CTLA4), and the tumor infiltrating lymphocytes (TILs) were TCRβ CDR3 sequenced on therapy without knowledge of the relevant recognized tumor antigens. In this dataset, the ground truth label corresponds not to an antigen-specificity but rather to the mode of therapy each mouse received. In the *CheckMate-038* dataset, patients with inoperable melanoma received either α-PD1 or α-PD1+α-CTLA4 and the ground truth label corresponds to radiographic response via RECIST v1.1 criterion. This latter analysis represents a test of DeepTCR as a pretreatment biomarker predictive of response in a clinical immunotherapy setting.

The main building block of all architectures in DeepTCR utilizes a common method of TCR featurization (Fig. 1a). First, any of the available α or β-chain CDR3 variable length sequences are provided to the network and are embedded via the use of a trainable embedding layer, as described by *Sidhom et.al* ^6^, to learn properties/features of the amino acids and transform the sequences from a discrete to continuous numerical space. Subsequently, a 3-layer CNN is used to extract sequence-based features from both chains. Additionally, V/D/J gene usage is provided to the network as a categorical variable in a one-hot representation. A trainable embedding layer is again leveraged to learn features of the V/D/J gene segments and transform them from a discrete to continuous numerical space. These features are then concatenated within the network to provide a joint representation of the sequence through its CDR3 sequences and V/D/J gene usage. All architectures implemented were created with Google’s TensorFlow™ deep learning library, allowing us to utilize Graphics Processing Units (GPU’s) for training and thus handle the large nature of TCR-Seq data in a time efficient and high-throughput manner.

**Figure 1.**
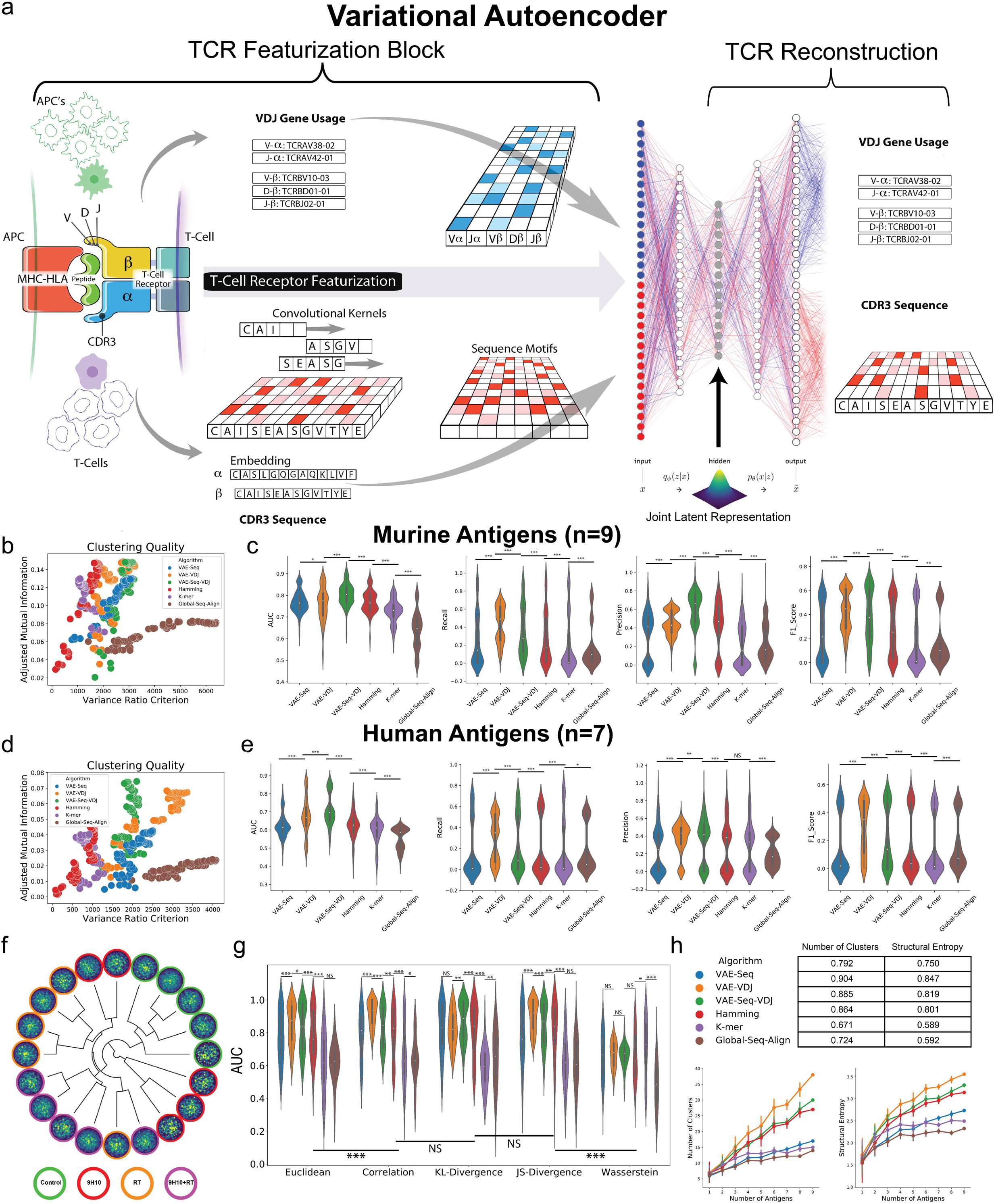
Unsupervised TCR Sequence & Repertoire Representation. **(a)** In order to represent a given T-Cell Receptor (TCR) we have implemented a variational autoencoder (VAE) to take the CDR3 sequences from both the α and β chains along with their corresponding V, D, and J-gene usage and learn a joint representation of these inputs. The CDR3 sequences serve as input to the encoder side of the network as discrete inputs with right zero-padding and a trainable embedding layer is used to transform the sequence from a discrete to a continuous numerical domain. Convolutional layers are then applied to learn sequence motifs from the CDR3 regions. As for the V/D/J gene inputs, these are represented as categorical variables which are then transformed with a trainable embedding layer to learn a continuous representation of the gene usage of a given TCR. Finally, these inputs are concatenated together passed through fully connected layers to result in the latent representation of the TCR. As is traditionally done in VAE networks, this latent space is parametrized by a multidimensional unit Gaussian distribution. This latent representation is then sampled from in order to reconstruct the input CDR3 sequences and V/D/J genes through the decoder side of the network. Finally, the weights of the neural network are trained via gradient descent to jointly minimize both the reconstruction and variational loss. The trained network is then used to take a given TCR and represent it in a continuous numerical domain for downstream analysis such as clustering. **(b,d)** In order to assess the quality of various methods of TCR featurization, we derived TCR distances from the various featurization methods (VAE-Seq, VAE-VDJ, VAE-Seq-VDJ, Hamming, K-mer, Global-Seq-Align) and applied an agglomerative clustering algorithm varying the number of clusters evenly from 5-100 and measured the variance ratio criterion of the clustering solutions and the adjusted mutual information from the clustering solutions to the ground truth antigen labels for both 9 murine and 7 human antigens. Featurization methods that encourage high quality clusters that capture a high degree of information of the label (i.e. antigen specificity) should have a high variance ratio criterion and high adjusted mutual information. **(c,e)** In order to benchmark the ability of various methods of TCR featurization to correctly classify a TCR sequence to its antigen, we applied a K-Nearest-Neighbors instance-based classification algorithm (varying K evenly from 1-500, Supplementary Fig. 3-10) to the derived TCR distances on the 9 murine and 7 human tetramer-sorted antigen-specific T-cells and assessed classification performance via 5-fold cross-validation strategy, measuring AUC, Recall, Precision, and F1 Score. Comparisons of performance were assessed at every value of K to determine statistical differences between methods. (Paired two-sample t-test, * : p < 0.05, ** : p < 0.01, *** : p < 0.001). **(f)** We developed a repertoire dendrogram to represent and compare TCR repertoires in an unsupervised fashion given their underlying TCR sequence features. We take featured TCR sequences from the VAE and applying a network-based clustering algorithm (PhenoGraph) to cluster the sequences in a sample-agnostic fashion. Every sample is then described by the proportion vector over the clustering solution. This vector is used to compute an inter-repertoire symmetric Kullback-Leibler (KL) divergence, which is then represented in the repertoire dendrogram using complete linkage. UMAP representation is learned from the featurized data to create a 2d representation of the data for each sample that is visualized at the leaves of the repertoire dendrogram, which is shown as a 2d histogram of the UMAP representation. When applied to TCR repertoires from TIL (*Rudqvist_2017),* the repertoire dendrogram is able to visualize and represent the relationships between the samples. **(g)** In order to assess the quality of using various TCR featurization methods to represent a given repertoire, we applied a K-Nearest Neighbors classifier (varying K evenly from 1-16, Supplementary Fig.12-15) to the proportional vector representing the distribution of a given TCR repertoire over a clustering solution of data taken from the *Rudqvist_2017* dataset. We employed a 5-fold cross-validation strategy and assessed classification performance via AUC as well as Recall, Precision, F1 Score (Supplementary Fig. 12-15) for how well different methods could classify a given repertoire to the therapy that mouse/sample received. We further assessed different distance metrics including Euclidean, Correlation, a symmetric KL-divergence, JS-divergence, and Wasserstein distance to base the KNN. (Paired two-sample t-test, * : p < 0.05, ** : p < 0.01, *** : p < 0.001) **(h)** To assess and quantify structural diversity of a TCR repertoire, we created *in-silico* repertoires by mixing between 1 and 9 murine antigen-specific populations and following featurization by all tested methods and clustering the given repertoire via PhenoGraph, we measured the number of clusters and Shannon’s entropy of the proportional vector over those clusters and measured the Pearson’s correlation coefficient of these metrics with the number of antigens that were mixed together to determine which featurization methods best captured the true antigenic diversity of the repertoire.

We first implemented this method of TCR featurization within the unsupervised learning setting in order to learn the underlying distribution of the sequence data in high-dimensional space for the purpose of 1) clustering TCR sequences that likely recognize the same antigen and 2) quantifying similarity between repertoires based on their structural composition. In order to learn the underlying distribution of the data in a structurally informed latent space, we implemented a variational autoencoder (VAE) as autoencoders have been previously described as a common dimensionality reduction/data re-representation technique^30,31^. Our implementation of a variational autoencoder (Fig. 1a) starts by taking a TCR sequence and following featurization as described previously, is transformed into a latent space that is parametrized by a multidimensional unit Gaussian distribution. The sequences and V/D/J gene inputs are then reconstructed from the latent space through the use of deconvolutional and fully-connected layers.

In order to assess the value of using deep learning as method of TCR featurization, we collected data for tetramer-sorted antigen-specific cells for 9 murine and 7 human antigens where the ground truth label corresponds to a particular antigen-specificity for an individual sequence^20–22^. We benchmarked the VAE against featurizations of TCR’s based on Hamming distances, K-mer representation, and global sequence alignment. Prior methods including *GLIPH* and *TCRdist* both use Hamming distances while *ImmunoMap* uses a global sequence alignment^20,22,27^. Our implementation of the Hamming distance method was directly benchmarked on the *Glanville_2017* dataset against the original *GLIPH* algorithm demonstrating improved clustering accuracy as measured in the original manuscript (Supplementary Fig. 2). For the VAE, we benchmarked the algorithm with different types of inputs to the network including just the β-chain CDR3 (VAE-Seq), just the V/D/J gene usage (VAE-VDJ), and the combination of both inputs (VAE-Seq-VDJ). First, to benchmark these various methods of featurization in clustering antigen-specific TCR’s, we ran an agglomerative clustering algorithm varying the number of clusters from 5 to 100 and then assessed the variance ratio criterion of the clustering solutions and the adjusted mutual information from the clustering solutions to the ground truth antigen labels (scikit-learn)^32,33^. We noted that the VAE methods maintained the highest variance ratio criterion while also maintaining a high adjusted mutual information to the ground truth labels for both murine and human datasets (Fig. 1b,d). To further query the value of these learned features in correctly clustering sequences of the same specificity, we applied a K-Nearest Neighbors Algorithm across a wide range of K-values using a 5-fold cross-validation strategy and assessed performance metrics of the classifier including AUC, Recall, Precision, and F1-Score for all featurization methods (Fig. 1c,e, Supplementary Fig. 3-10)^34^. We noted that across all performance metrics, the VAE methods outperformed current state-of-the-art approaches for TCR featurization. Furthermore, using both sequence and V/D/J gene usage resulted in the highest AUC performance for both the murine and human antigens suggesting both types of inputs provide distinct and contributary information to antigen-specificity assignment in addition to encouraging a featurization of the TCR that is length invariant (Supplementary Fig. 11).

While most of the previous work has focused on analyzing TCR-Seq at a single sequence level, investigators often desire to do repertoire-level comparisons and analyses. In light of this need, and as others have attemped^35,36^, we sought to create a downstream pipeline that could use this improved featurization to make inter-repertoire comparisons. Our approach is driven by the VAE TCR featurizations described above, in which a network graph-based clustering algorithm (PhenoGraph) is applied to result in a proportionality vector of the various TCR sequence clusters observed within a given repertoire^37,38^. To make inter-repertoire comparisons we measure the distance between these proportionality vectors using variety of distance metrics including Euclidean, Correlation, a symmetric KL-divergence, JS-divergence, and Wasserstein distance. To visualize these data, we developed a Repertoire Dendrogram that shows each repertoire in a 2d UMAP representation and the inter-repertoire relationships through a dendrogram determined by the aforementioned distance metrics (Fig. 1f)^39^. When applying a K-Nearest Neighbor classifier on the proportionality vectors to predict the treatment label of these murine repertoires in the *Rudqvist_2017* dataset we found that: (a) Correlation, KL-divergence, and JS-divergence outperformed Euclidean and Wasserstein distance metrics (b) VAE methods outperformed the state-of-the-art featurization methods (Fig. 1g, Supplementary Fig. 12-15). Furthermore, using simulated data comprised of 1-9 previously described murine antigens, we showed that that the VAE-VDJ, VAE-VDJ-Seq, and Hamming methods were the top performing methods in describing the expected antigenic diversity of these *in-silico* repertoires (Fig. 1h).

To leverage ground truth labels often associated with TCR-Seq, we developed a fully supervised model, that learns sequence specific motifs to correctly classify sequences by their labels (Fig. 2a), in which we observed that our supervised approach improved performance over the previously described unsupervised VAE approach (Fig. 2b, Supplementary Fig. 16) and a more conventional Random Forest (RF) & Support Vector Machine (SVM) (Supplementary Fig. 17)^40,41^. In addition, being able to extract knowledge from the network can inform relevant motifs for antigen-specific recognition. Therefore, we established a method by which we could identify the most predictive sequences for a given class and query the learned kernels/motifs these sequences were uniquely activating (Fig. 2c).

**Figure 2.**
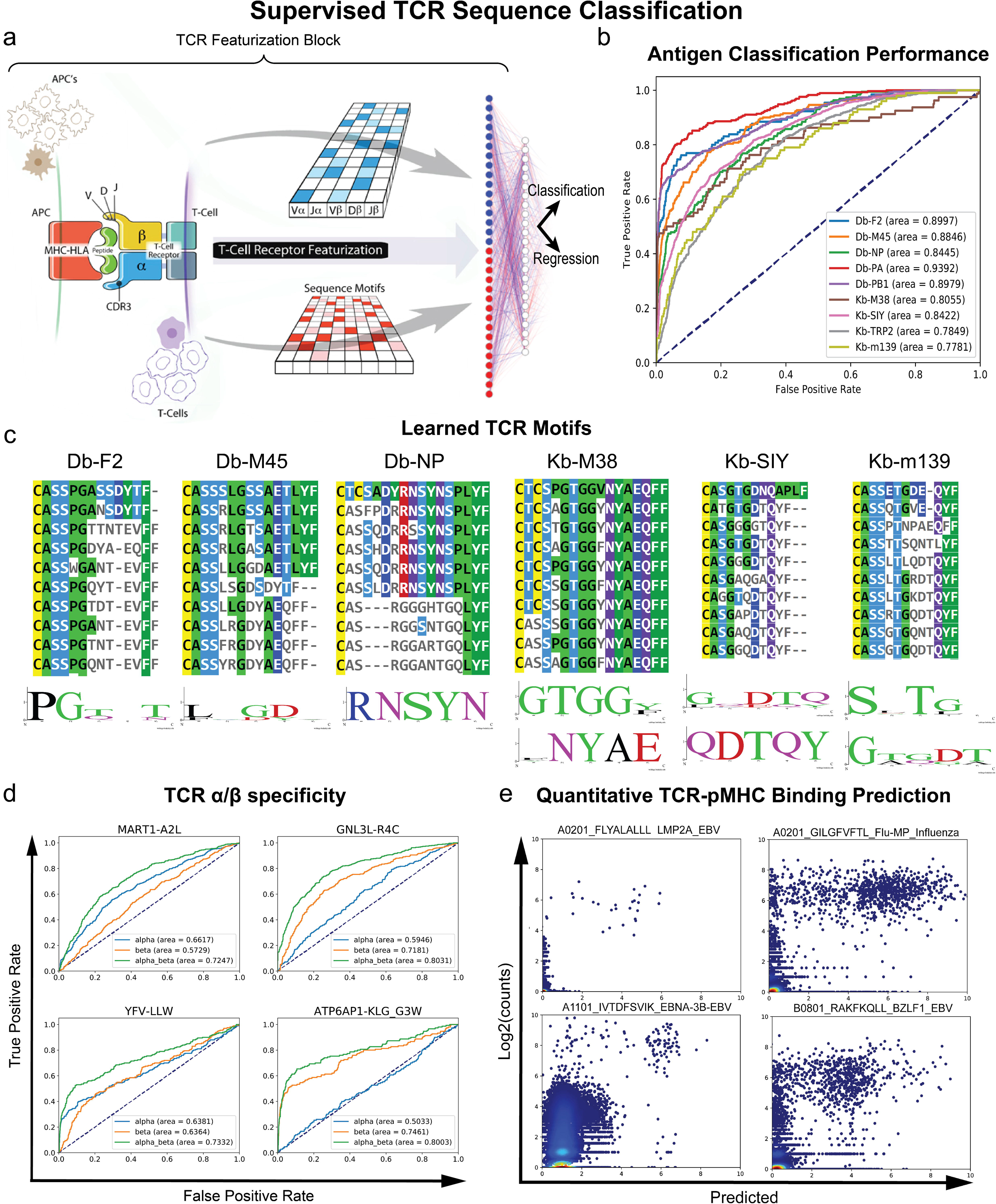
Supervised TCR Sequence Representation. **(a)** Network Architecture Schema: Previously described TCR Featurization block is implemented to featurize a TCR sequence and then either output a label (i.e. antigen-specificity) or continuous regressed value (i.e. affinity measurement). **(b)** Supervised TCR Sequence classifier was trained/tested on 9 murine antigen-specific TCR sequences via 10-fold Monte-Carlo cross-validation strategy where classification performance, assessed via AUC measurements, was measured on the test sets. Classification performance was also benchmarked against DeepTCR’s unsupervised instance-based learning classifier and classical Random Forest (RF) and Support Vector Machine (SVM) algorithms (Supplementary Fig. 16, 17). **(c)** Representative Db and Kb murine antigens where top predicted CDR3 sequences are shown via multiple-sequence alignment and learned kernels for these representative sequences are visualized below the alignment. **(d)** In order to assess the contribution of the TCR α and β-chains in defining antigen-specificity, T-cells from healthy individuals were sorted via DNA-barcoded antigen tetramers for either public (Number of Unique α/β pairings: MART1-A2L = 72, YFV-LLW = 32) or cancer neoantigens (Number of Unique α/β pairings: GNL3L-R4C = 58, ATP6AP1-KLG_G3W = 24) from the literature and were sent for single-cell sequencing of the α/β chains (*Zhang_2018)*. DeepTCR’s sequence classifier was then used to assess predictive structural signature using input of α alone, β alone, or combined to predict antigen specificity as characterized by AUC from ROC analysis. **(e)** In order to test the ability of a supervised deep learning method to learn and regress continuous value outputs, we collected published single-cell data from 10x Genomics where 57,229 unique α/β pairs were collected with a count-based measurement (as a proxy for binding affinity) to 44 specific peptide-MHC (pMHC) multimers and 6 negative controls. A 5-fold cross-validation strategy was employed on every antigen to obtain independently predicted regression values for every α/β pair to a given antigen and predicted vs. actual counts are shown for a select 4 antigens.

One of the key advantages of DeepTCR is its ability to learn a joint representation of a TCR through any of the aforementioned inputs of the α and β chain as compared to a separately modeled process^27^. To highlight our ability to model possible interactions of the α and β chain, we applied DeepTCR to recent single-cell data, *Zhang_2018*, in which coupled α and β chain sequences are available^28^. DeepTCR was able to reveal the contribution of each chain to specificity for the antigen and the level of synergy in a quantitative fashion (Fig. 2d), demonstrating that information from both TCR chains are important for predicting antigen-specificity across both common and mutation-associated neoantigens (MANAs). The implications of this finding in cancer immunology are particularly relevant as the field commonly only sequences the β-chain when assessing the adaptive immune response to tumors^8,11,12,15,29,42^. Finally, the binding of a TCR to a peptide-Major Histocompatibility Complex (pMHC) is not usually considered a binary phenomenon but rather one that is characterized by a binding affinity. Therefore, we applied DeepTCR to regress UMI (unique molecular identifier) counts as a proxy for binding affinity (a caveat to this assumption being that differing TCR expression levels can also affect the UMI counts) as available in a second single-cell data set published by 10x Genomics where the affinity of 57,229 unique α/β pairs to 44 specific pMHC multimers and 6 negative controls was characterized. DeepTCR was able to identify TCRs that have both high observed UMI counts and a predictive structural signature, providing a tool to better isolate antigen-specific TCRs (Fig. 2e).

Building on the supervised sequence classifier, we then wanted to design an architecture that could learn from a label applied to a whole repertoire of TCR sequences. This type of problem can be poised as ‘weakly supervised’ as the repertoire label may only apply to a subset of the sequences^43^. Our supervised repertoire classifier was formulated as a supervised multi-instance learning (MIL) algorithm that is able to extract meaningful concepts that may lie within large repertoires of many sequences through the use of a multi-head attention (Fig. 3a). We applied our DeepTCR MIL repertoire classifier to the *Rudqvist_2017* dataset, in which a significant increase in performance was observed over the previously described sequence level classifier (Fig. 3b). Interestingly, the post treatment repertoire signature was the least predictive in the cohorts of mice that received checkpoint block inhibition (α-CTLA4), compatible with the notion that α-CTLA4 was acting by shaping the structural diversity of the repertoire towards immune recognition of personalized neoantigens, as has been suggested in prior work as a possible mechanism of action^44,45^. Finally, not only are supervised approaches valuable for their predictive utility but they can also provide a different and more informative featurization and description of the TCR repertoire guided by ground truth labels (Fig. 3c).

**Figure 3.**
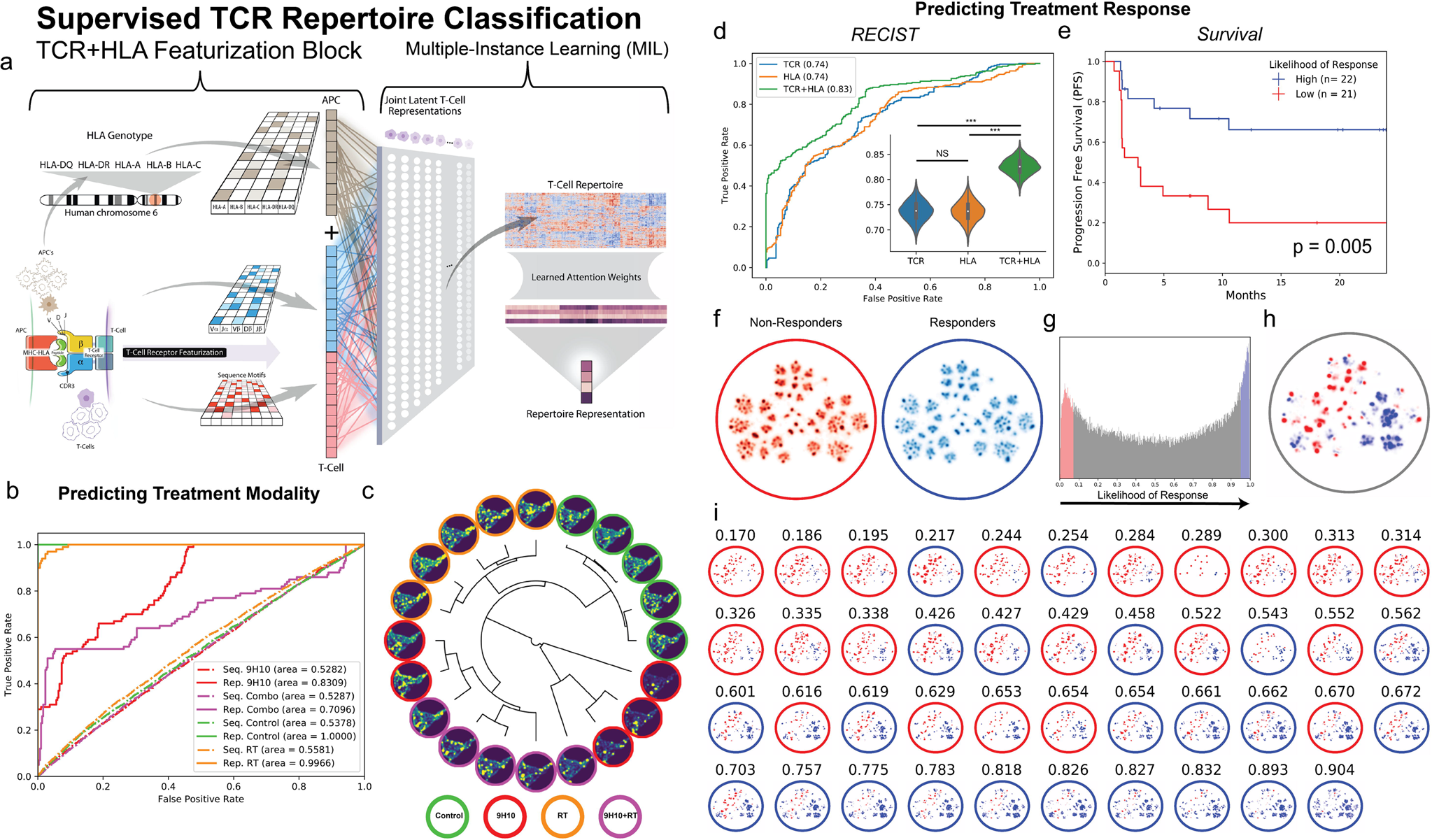
Supervised TCR Repertoire Representation. **(a)** Multiple-Instance Learner (MIL) for classifying a TCR repertoire. First, we expand the TCR Featurization block to incorporate the HLA background within which a given collection of TCR’s were observed within. The HLA background of a sample/individual is provided to the neural network in a multi-hot representation that is re-represented in a learned continuous embedding layer and concatenated to the continuous learned representation of the TCR. Second, we implement a multi-head attention mechanism to make sequence assignments to concepts within the sample. The number of concepts in the model is a hyperparameter, which can be varied by the user depending on the heterogeneity expected in the repertoires. Of note, this assignment of a sequence to a concept is done through an adaptive activation function that outputs a value between 0 and 1, allowing the network to put attention on the sequences that are relevant to the learning task. When taking the average of these assignments over all the cells in a repertoire, this results in a value within the neural network that directly corresponds to the proportion of the repertoire that is described by that learned concept. These proportions of concepts in the repertoire are then sent into a final traditional classification layer. **(b)** The DeepTCR supervised sequence classifier was first applied to sequences from *Rudqvist_2017* dataset to assess the predictive signature to correctly classify a given TCR sequence to the therapy of the mouse it was observed within (5-fold cross-validation). The supervised MIL repertoire classifier was then applied to samples from the *Rudqvist_2017* dataset to assess the predictive signature to correctly classify a collection/repertoire of TCR sequences to the therapy of the mouse it was observed within (100 Monte-Carlo simulations with train size: 16, test size: 4). Performance was assessed through measuring the AUC of the classifiers (sequence classifier – dotted lines, repertoire classifier – solid lines). **(c)** TCR sequence level predictions from the repertoire classifier trained on the *Rudqvist_2017* dataset were taken and averaged over the 100 MC simulations to create a supervised featurization of the TCR repertoires on which a repertoire dendrogram was used to display the relationship between mice and the distribution of TCR sequences in each mouse. **(d)** Pre-treatment tumor biopsies were collected and TCR-Seq was performed from 43 patients enrolled in the *CheckMate-038* (parts 2-4) clinical trial where they were either treated with α-PD1 monotherapy (9 patients) or α-PD1+ α-CTLA combination therapy (34 patients) and followed for radiographic response to therapy via RECIST v1.1 (Supplementary Fig. 18). Complete Responders and Partial Responders (CRPR) were denoted as responders to therapy while Stable Disease and Progressive Disease (SDPD) were denoted as non-responders to therapy. Receiver Operating Characteristics (ROC) Curves were created for predicting response (complete response, partial response) to immunotherapy given either TCR, HLA, or TCR+HLA information to the supervised repertoire classifier (100 Monte-Carlo simulations with train size: 37, test size: 6). Bootstrap analyses (5000 iterations) were performed to construct confidence intervals (CI) around AUC values and assess differences in model performance, in which each AUC per sampling was compared in a paired manner across the three models designed above. The null hypothesis of two models exhibiting equivalent performance was rejected if the bootstrap difference did not cross 0. (*** : 99.9% CI). **(e)** The likelihood of response generated by the TCR+HLA model was dichotomized into “High” and “Low” using the median predicted value in this cohort (taken over the MC test sets and averaged per sample) and the Kaplan-Meier (KM) curves were shown for progression free survival (PFS), log-rank p-value = 0.005. **(f)** In order to provide a descriptive understanding of the T-cell response in responders and non-responders in the *CheckMate-038* clinical trial, we sought to characterize the distribution of the TCR repertoire in this cohort of patients. Data from *CheckMate-038* were used to train a VAE on all sequence data (incorporating TCR+HLA information) in a sample and class agnostic fashion. The distribution of responders and non-responders repertoires were visualized via UMAP of the unsupervised VAE featurization (per sample distribution shown in Supplementary Fig. 23). **(g)** In order to visualize the distribution of the highly predictive TCR sequences, a per-sequence prediction value was assessed following each MC simulation on the TCR’s within the independent test set, assigning the probability that a given TCR had a responder signature. Over the 100 MC simulations, each sequence in the cohort is assigned multiple prediction values that are averaged over all simulations to serve as a consensus predicted value for each sequence in this cohort of patients. **(h)** Top 10% of sequences in responders and non-responders were selected and visualized over the entire cohort and on a per-sample basis **(i)** where edge color denotes the ground truth label of the sample (non-responder = red, responder = blue) and average predicted likelihood taken over MC simulations to respond to treatment shown above each patient’s distribution.

Finally, we applied DeepTCR to pre-treatment tissues in a clinically relevant dataset, *CheckMate-038* (Supplementary Fig. 18). As compared to the *Rudqvist_2017* dataset from genetically identical mice, in the human setting, individuals have different HLA (Human Leukocyte Antigen) backgrounds and their tumors have unique mutation-associated antigens and tumor-associated antigens; thus, TCR’s that are structurally homologous would likely not be recognizing the same tumor antigens. In order to account for this complexity in human data, we encoded HLA information into the featurization architecture that could be used to fuse HLA information into a joint TCR-HLA antigenic latent space (Fig. 3a).

We observed that when applying DeepTCR to predict response to immunotherapy, the joint representation of TCR with HLA outperformed models of DeepTCR that used TCR sequence or HLA genotype information alone (AUCs; TCR = 0.74, HLA = 0.74, TCR+HLA = 0.83, random permutation testing = 0.515; Fig. 3d, stratified by treatment cohort in Supplementary Fig. 19). The resulting DeepTCR’s TCR+HLA model predictions of likelihood to respond to treatment also significantly stratified progression free survival (PFS) in this cohort of patients (Fig. 3e). Interestingly, the TCR+HLA model exhibited less variability during cross validation assessments as compared to either the TCR or HLA model alone (Supplementary Fig. 20); and it appears that the addition of HLA genotype conditions the model to learn different information from the TCR sequence data (Supplementary Fig. 21). We also found that in this cohort, the DeepTCR TCR+HLA model performed comparably to conventional biomarkers that have been previously described to stratify response in immunotherapy-treated patients (AUCs; PD-L1 (IHC TPS score) = 0.691, TMB (exome-based) = 0.773, TCR Clonality = 0.635, Total T-cell count = 0.820; Supplementary Fig. 22). And while similar performance (based on AUC) is observed with total T-cell counts (a surrogate of the degree of T-cell tumor infiltration) and DeepTCR, these predictors are independent on multivariate logistic regression analyses and as such plausibly represent complementary information. Further characterization of multivariate models of response prediction should be described and validated in conjunction with separate, independent cohorts. While an independent validation cohort is not yet available, all performance assessments reported herein for DeepTCR were based on rigorous Monto-Carlo cross validation assessments with a strict adherence to partitioning of data points used to train the model vs data points used to evaluate trained model performance.

Finally, to characterize the distribution of the TCR repertoires in patients who either responded or did not respond to treatment, we trained a VAE on all data to obtain an unsupervised featurization to visualize (via UMAP) the distribution of non-responders and responders (Fig. 3f, per sample distributions shown in Supplementary Fig. 23). While no difference in the overall distributions can be appreciated when looking at all the data, we found that when filtering the TCR sequences to the top and bottom 10% predictive sequences (Fig. 3g), we were able to visualize differences in the TCR repertoires between responders and non-responders (Fig. 3h), illustrating the functionality of the MIL algorithm to ‘denoise’ TCR repertoire. We noted that not only are the distributions within responders and non-responders multi-modal, but these multiple modes are shared between patients (Fig. 3i). These findings suggest that the antigenic response is not only different between responders and non-responders to checkpoint blockade but each cohort of patients contains a set of T-cell responses that likely recognize a broad class of structurally-related antigens.

NGS has become one of the largest sources of big data in the biological sciences, and deep learning is a promising modality for analyzing big data where features need to be learned. In this work, we present DeepTCR, a collection of unsupervised and supervised deep learning approaches to characterize TCR-Seq data for both descriptive and predictive purposes. We first demonstrate that by using a variational autoencoder to do unsupervised learning with an improved method of TCR featurization, we can better relate and compare samples at the sequence and repertoire level. More significantly, we develop supervised methods in applications where labels can greatly help the learning process, such as when there is buried signal in a large sample of sequences. Finally, we introduce a method by which to re-represent TCR information in the context of HLA background and demonstrate this approach to predict response to immunotherapy in a clinical trial, despite the intrinsic variation in HLA type, tumor antigens, cancer genetics and epigenetics among patients. Given the ongoing mandate in personalized immunotherapy for improved predictive biomarkers of therapeutic response, DeepTCR may have near term value in clinical oncology.

## Methods

### Data Curation

TCR sequencing files were collected as raw tsv/csv formatted files (Supplementary Fig. 1) from the various sources cited within the manuscript. Sequencing files were parsed to take the amino acid sequence of the CDR3 after removing unproductive sequences. Clones with different nucleotide sequences but the same amino acid sequence were aggregated together under one amino acid sequence and their reads were summed to determine their relative abundance. Within the parsing code, we additionally specified to ignore sequences that used non-IUPAC letters (*,X,O) and removed sequences that were greater than 40 amino acids in length. For the purpose of the algorithm, the maximum length can be altered but we chose 40 as we did not expect any real sequences to be longer than this length.

### CheckMate-038 Experimental Model and Subject Details

CheckMate-038 is a multi-arm, multi-institutional, institutional-review-board-approved, prospective study (CA209-038; NCT01621490). Patients in Parts 2-4 received either 3 mg/kg nivolumab every 2 weeks (N = 21), or 1 mg/kg nivolumab + 3 mg/kg ipilimumab every 3 weeks x 4 doses and then 3 mg/kg nivolumab every 2 weeks (N = 62), until progression or for a maximum of 2 years. Radiographic assessment of response was performed approximately every 8 weeks until progression. Progression was confirmed with a repeat CT scan typically 4 weeks later. Tumor response for patients was defined by RECIST v1.1. Response to therapy indicates best overall response unless otherwise indicated. All patients underwent biopsy of a metastatic lesion before commencing therapy (1–7 days before first dose of therapy). Tumor tissue was divided for formalin-fixation, paraffin-embedding (FFPE) or storage with RNAlater (Ambion) for subsequent RNA/DNA extraction. PD-L1 expression on the tumor cell surface was assessed in FFPE samples at a central laboratory (Dako 28-8 antibody).

### CheckMate-038 TCR-Seq & HLA Data Generation

Tumor biopsy samples were collected prior to initiation of therapy and stored in RNAlater. DNA was extracted and submitted to Adaptive Biotechnologies for survey level TCR β-chain sequencing, where targeted amplicon libraries were prepared by multiplex PCR targeting all TCR β-chain V and J gene segments and sequenced using the Illumina HiSeq system^46,47^. Data for individual TCR sequences, including V and J gene segment identification and CDR3 sequence, were obtained from Adaptive Biotechnologies for analysis via DeepTCR. Tumor biopsy DNA was also sent for whole-exome sequencing (Personal Genome Diagnostics) to determine Total Mutational Burden (TMB) and OptiType was utilized to infer HLA genotype of the patients^48^. Data from patients who consented to deposition will be submitted to the European Genome-phenome Archive.

### Data Transformations

In order to allow a neural network to train from sequence data, we converted the amino acids to numbers between 0-19 representing the 20 possible amino acids. These were then one-hot encoded as to provide a categorical and discrete representation of the amino acids in numerical space. This process was applied prior to all networks being trained. For analyses where V/D/J gene usage or MHC-HLA genes were included, these genes were represented as categorical variables and one-hot (V/D/J) or multi-hot (MHC-HLA) encoded as inputs for the neural network.

### TCR Featurization Block

The core of all deep learning architectures is the TCR Featurization Block which takes the various sequence data for a given TCR and transforms it to a latent joint representation of all its inputs. For the α/β CDR3 sequences, we take variable length right-padded sequence data which has been encoded in one-hot representation and first apply an embedding layer which transforms this one-hot representation to a trainable continuous representation of dimensionality 64. This embedding layer learns features of each amino acid allowing the network to learn amino acids which may play similar roles in antigen-binding in the context of the TCR. Following this transformation, 3 convolutional layers are applied to the continuous representation of the CDR3 sequences. The kernel, stride sizes, and number of feature maps were (kernel: 5, stride: 1, feature maps: 32), (kernel: 3, stride: 3, feature maps: 64), (kernel: 3, stride: 3, feature maps: 128) respectively for the 3 layers. If the convolutional stack is being used within the variational autoencoder, the output of the final convolutional layer is flattened. If the convolutional stack is being used within either the supervised sequence classifier or repertoire classifier, the global max pooling operation is applied across the length of the sequence to provide the ability for the network to learn length-invariant motifs.

If V/D/J gene information is provided as an input to the network, this data is represented first as categorical variable with a one-hot encoding to the network. Once again, we apply a trainable embedding layer which transforms this one-hot representation to a continuous representation of dimensionality 48. This transformation produces the featurization of the V/D/J genes. Finally, if MHC-HLA information is provided (i.e. the background the TCR was observed within), the data is represented as a multi-label categorical variable with a multi-hot encoding, allowing a description of the HLA background of an individual as a combination of multiple alleles. A trainable embedding layer is applied which transforms this multi-hot representation to a continuous representation of dimensionality 12. This transformation produces the featurization of the MHC-HLA background of an individual. This HLA representation is then concatenated to the TCR features of a given receptor and one fully connected layer is always implemented at this point to generate a new set of features that is jointly learned between the HLA background and the TCR features. This representation can be thought of as learning the peptide-antigen that would be recognized via this combination of TCR and HLA molecule.

After all inputs to the network have been featurized, they are concatenated and this completes the TCR Featurization Block where a TCR is described by a vector of continuous variables that describe all of the possible CDR3 sequences, V/D/J gene usage, and MHC-HLA background or context within which a TCR was observed within. This TCR Featurization Block is used as the main building block for all networks described and used in the manuscript.

### Training VAE

In order to train the VAE, following creation of the computational graph as described in the manuscript and main figure, we applied an Adam Optimizer (learning rate = 0.001) to minimize a reconstruction loss and a variational loss. The reconstruction loss is the cross-entropy loss between the reconstructed sequence (S) and the one-hot encoded tensor of the input sequence (L) across the *i*th position in the sequence (1). The variational loss is the Kullback–Leibler (KL) divergence between the distributions of the latent variables and a unit gaussian (2).

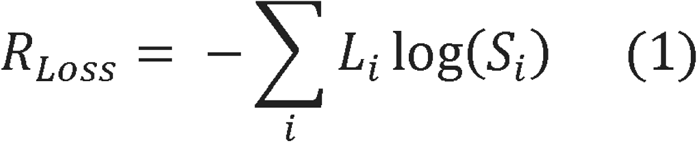

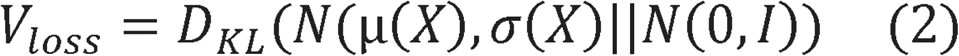

The variational loss serves as a regularizer to the network as it prevents overfitting of the network and direct memorization of sequence to latent space and allows for meaningful downstream clustering of the sequences in their latent representation^30,31^. The variational autoencoder was trained until convergence criteria were met. Features for all sequences were then extracted from the latent space and used for downstream analyses.

### Quantifying TCR Distance

In order to quantify the distance between TCR sequences from the latent representations produced by the variational autoencoder, we computed a Euclidean distance in this space to measure the distance between any two TCR sequences. For the K-mer representation, we also used a Euclidean distance on the K-mer count vector to measure the distance between any two TCR sequences. To compute Hamming distance, we used the scipy.pdist function on the integer representation of the sequences. For the global sequence alignment-based distance, we computed a symmetric distance based as previously described by *Sidhom et. al* in the ImmunoMap algorithm. Global sequence alignment was computed with BioPython’s pairwise2.align.globalxx functionality^20^.

### Clustering Antigen-Specific TCR Sequences

To assess the quality of the various featurization methods described in the study, we first applied an agglomerative clustering algorithm (scikit-learn) to the previously described TCR distances from the various VAE methods along with the Hamming, K-mer, and Global Sequence Alignment distance metrics. We varied the number of clusters for the algorithm from 5-100 clusters and measured the Variance Ratio Criterion (Calinski and Harabasz score) as well as the Adjusted Mutual Information across all the clustering solutions for all the described featurization methods^32,33^. By choosing these two metrics to quantify the robustness of the clustering solutions on the latent features, we first assessed the ratio of the within-cluster dispersion to the between-cluster dispersion as a measure for the ‘compactness’ of the clustering solution via the Variance Ratio Criterion and then quantified using information theoretic principles to quantitate how much of the information about the antigen-specificity was being captured by the clustering solution. Methods that provide a feature space optimal for applying clustering algorithms would have a high Variance Ratio Criterion as well as a high Adjusted Mutual Information. These metrics were applied to both the murine and human antigen-specific TCR’s to compare the various featurization methods.

### Training K-Nearest Neighbor (KNN) Algorithm on TCR Sequences

In order to assess the quality of the various featurization methods describes in the study, we also applied a KNN on to the previously described TCR distances derived from the various VAE methods along with the Hamming, K-mer, and Global Sequence Alignment distance metrics. We employed a 5-fold cross-validation strategy to split the data and then assessed performance on the left-out fold of the data. Furthermore, we varied the value of K in the KNN evenly from 1-500 in order to further assess the robustness of the featurization/KNN across a wide variety of K values. We were then able to do paired statistical analyses at each value of K to assess which featurization methods allowed for best downstream performance of the classifier to assign a TCR sequence to its correct cognate antigen.

### Repertoire Dendrogram

In order to visualize and compare TCR repertoires in an unsupervised fashion, we developed a visualization technique we termed a ‘Repertoire Dendrogram’ in which we utilize a dendrogram to display the relationships between separate TCR repertoires. In order to construct this dendrogram, we begin by applying a network graph-based clustering algorithm (PhenoGraph) on all the TCR sequences in a sample/repertoire agnostic fashion to learn naturally occurring clusters of concepts within the data^37^. We then determine the proportional distribution of all samples/repertoires across this clustering solution to describe each repertoire as its distribution across the clustering solution. We then measure the distance between samples/repertoires via taking some distance metric (i.e. KL divergence) between these proportional vectors and use this distance measurement to construct the dendrogram tree. Finally, we learn a 2d representation of the data through a UMAP transformation that allows us to display the sequences of a given repertoire in the leaf of each node on the tree. In this way, one can visualize which repertoires are similar in their TCR composition and what areas in this learned high-dimensional latent space are similar between repertoires^39^.

### Training K-Nearest Neighbor (KNN) Algorithm on TCR Repertoires

In order to assess the quality of the various featurization methods describes in the study to describe whole TCR repertoires, we utilized a K-Nearest Neighbor approach to classify TCR repertoires based on their proportional vectors of a network graph-based clustering solution (described above). Following clustering via PhenoGraph across all tested featurization methods, computing the proportional vectors, and computed distances using a variety of distance metrics (i.e. Euclidean, Correlation, a symmetric KL-divergence, JS-divergence, and Wasserstein distance) we trained/tested a KNN algorithm to predict the treatment label from the *Rudqvist_2017* dataset^29^. We employed a 5-fold cross-validation strategy and varied the K-parameter in the KNN from 1-16 where we assessed the predictive power on the left-out fold. We made pairwise statistical comparisons at all values of K to assess the performance of various featurization methods as well as various distance metrics applied to the proportional vector derived from these aforementioned featurization methods.

### Measuring Antigenic Diversity in TCR Repertoire

As has been noted, TCR-Seq is often an assay performed on a collection or repertoire of T-cells. As is such in a biological function of a repertoire, such as in cancer, this collection of T-cells may often have many antigen-specificities and an investigator my want to know the antigenic diversity of a TCR repertoire. We wanted to see whether a deep learning featurization could capture and describe the level of antigenic diversity in a TCR repertoire so we constructed simulated data sets *in-silico* where we mixed TCR sequences with known antigen-specificities from anywhere between 1-9 murine antigens, applied DeepTCR’s VAE to featurize the sequences with PhenoGraph clustering to derive proportional vectors of the distribution of each simulated repertoire. We then measured the number of clusters that were formed and the Shannon’s entropy of this distribution and used these as measured for the structural or antigenic diversity of a given TCR repertoire.

### Training Receptor Classifier

In order to train the receptor classifier, we use the TCR Featurization Block as described previously to featurize the input data which can include any or all of α/β CDR3, the V/D/J gene usage, and MHC-HLA input. The main difference in this featurization is that for the CDR3 sequences, we employ a global max pooling operation after the final convolutional layer to allow for translational invariance of motifs within the CDR3 sequence. The final feature space is directly sent to a classification layer where the number of final nodes is equivalent to the number of classes. In the case of a regression task where the receptor is being regressed to a continuous label, the feature space is sent to a single node. In the case of a classification task, the network is trained using an Adam Optimizer (learning rate = 0.001) to minimize the cross-entropy loss between the softmaxed logits and the one-hot encoded representation of the discrete categorical outputs of the network. In the case of a regression task, the network is trained to minimize the mean squared error loss between the output of the final node in the network at the continuous label. Training was conducted by using 75% of the data for the training set, and 25% for validation and testing. The validation group of sequences was used to implement an early stopping algorithm.

### Training Repertoire Classifier

Designing an architecture for whole sample multi-instance classification presented unique challenges that were specific to the way TCR-Seq data is generated. Following featurization via the described TCR Featurization Block, we needed an architecture that could handle applying a label to a collection of these featurized sequences. In order to solve this multi-instance problem, we developed a multi-head attention mechanism that uses an adaptive activation function to make an assignment for each TCR sequence to a learned concept within the data. In order to design this activation function, we based it on the inverse square root unit (ISRU) function which is an algebraic form of the sigmoid function. While activation functions in neural networks are often fixed and have no trainable parameters (i.e. relu, sigmoid), we noted difficulty in training a sigmoid function to make an assignment between 0 and 1 due to the commonly cited problem of diminishing gradients with the use of sigmoid activation functions. By creating an adaptive ISRU function with a trainable α and β parameter *(3),* we found this improved the training of our network and allowed us to make a sequence-level assignment between 0 and 1 for each sequence to each learned concept in the model.

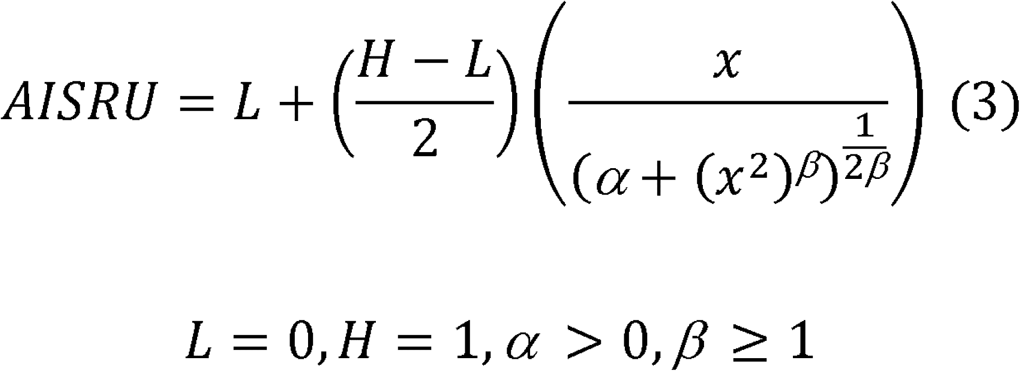

This average of these assignments is taken over the sample to come up with what can be interpreted as the proportion of the repertoire that contains the learned concept. This vector of proportion features is then fed directly into the classification layer. The network is trained with an Adam Optimizer (learning rate = 0.001) to minimize the cross-entropy loss between the softmaxed logits and the one-hot encoded representation of the discrete categorical outputs of the network.

Due to the small nature of the *Rudqvist_2017* and *CheckMate-038* datasets, training had to be accomplished in a way to prevent over-fitting of the repertoire classifier. Therefore, to train the repertoire classifier on these datasets, we employed a Monte-Carlo (MC) cross-validation where in the hinge-loss was used during model training, which prevents the model from further reducing the loss of any given sample below a defined threshold. The idea behind this type of objective function is that once a sample has been predicted sufficiently correct, the network is not encouraged to drop its loss any further and thus, reduces over-fitting on the training data. Model training with this hinge-loss was stopped once a pre-defined thresholds was meet and in keeping Monte-Carlo (MC) cross-validation, model perform was assessed on the testing data of that train/test split. We then used a bootstrapping method where we sampled the MC predictions with replacement 5000 times to approximate a confidence interval around the AUC.

### Motif Identification

Neural networks are often treated as ‘black boxes’ where their value is largely in their predictive performance and not in understanding how the neural network is accomplishing its task. However, in the area of the biological sciences, there is not only the desire to create predictive tools but use these tools to inform our own understanding of the mechanisms at play. This area of research is often termed as improving the ‘explainability’ of neural networks. In biological sequence analytics such as DeepTCR, investigators want to be able to extract the features/motifs the neural network learned to accomplish its task. For the supervised learning architectures, we were able to identify motifs the network had learned by extracting the indices of where the kernels were activated following the global max pooling layer. The result of this operation is the network not only extracts the maximum value of a kernel over the length of the sequence but also deduces its position within the sequence. This can be then used to not only pick up which features are activated on a given sequence but where in the sequence this activation occurs, allowing us to identify the motifs that any given neuron in the net is learning. Sequence logos were created with https://weblogo.berkeley.edu/logo.cgi.

### Statistical Tests & Machine Learning Models

All statistical tests applied to data were implemented with the scipy.stats module. Classical machine learning techniques and performance metrics were implemented with scikit-learn.

## Code and Data availability

DeepTCR was written using Google’s TensorFlow™ deep learning library (https://github.com/tensorflow/tensorflow) and is available as a python package. Source code, comprehensive documentation, use-case tutorials, and all ancillary code (including all deep learning hyperparameters) to recreate all the figures in the text along with all data and results presented in this manuscript can be found at https://github.com/sidhomj/DeepTCR. DeepTCR can either be installed directly from Github or from PyPI at https://pypi.org/project/DeepTCR/.

## Acknowledgements

The authors thank the Bristol-Myers Squibb for funding the CheckMate-038 clinical trial and all associated studies and data generation presented in this manuscript along with the MARC/SU2C Foundation for providing financial support for the work of developing algorithmic pipelines presented in this manuscript. We would like to thank Suzanne L. Topalian for helpful discussions and critical review of the manuscript, James R. White for editorial assistance and reviewing the DeepTCR codebase, and Valsamo Anagnostou and Victor Velculescu for generating HLA haplotype data from WES data collected in this study. Finally, we would like to thank the patients and their families for their participation in this study.

