## Supplementary Material for "DeepTCR: a deep learning framework for understanding T-cell receptor sequence signatures within complex T-cell repertoires"

1. **Dataset Descriptions**
2. **Comparing clustering performance of Hamming Distance v. GLIPH**
3. **AUC Scores for K-Nearest Neighbors on Murine Antigens**
4. **Recall Scores for K-Nearest Neighbors on Murine Antigens**
5. **Precision Scores for K-Nearest Neighbors on Murine Antigens**
6. **F1 Scores for K-Nearest Neighbors on Murine Antigens**
7. **AUC Scores for K-Nearest Neighbors on Human Antigens**
8. **Recall Scores for K-Nearest Neighbors on Human Antigens**
9. **Precision Scores for K-Nearest Neighbors on Human Antigens**
10. **F1 Scores for K-Nearest Neighbors on Human Antigens**
11. **Assessing correlation between various distance metrics and length of the sequence**
12. **AUC Scores for K-Nearest Neighbors on samples taken from murine tumor-infiltrating lymphocytes (TIL)**
13. **Recall Scores for K-Nearest Neighbors on samples taken from murine tumor-infiltrating lymphocytes (TIL)**
14. **Precision Scores for K-Nearest Neighbors on samples taken from murine tumor-infiltrating lymphocytes (TIL)**
15. **F1 Scores for K-Nearest Neighbors on samples taken from murine tumor-infiltrating lymphocytes (TIL)**
16. **Benchmarking Classification Performance between unsupervised and supervised deep learning methods.**
17. **Benchmarking various machine learning methods to classify TCR sequences by their antigen-specificity**
18. **CheckMate-038 Clinical Trial Schema**
19. **DeepTCR Performance on CheckMate-038 data stratified by treatment cohort**
20. **Distribution of CheckMate-038 Sample Predictions from Various Models**
21. **Sample Predictions for CheckMate-038 trial patients from different repertoire classification models**
22. **Comparison of DeepTCR against other biomarkers in CheckMate-038**
23. **UMAP representation of unsupervised VAE featurization for all sequence data in CheckMate-038**

| Dataset | Host | Pathology | Description |
|---|---|---|---|
| Glanville_2017 | Human | Infectious Disease | T-cells were sorted and sequenced from peripheral blood mononuclear cells (PBMCs) from healthy donors for 7 Class-I specificities. |
| Sidhom_2017 | Murine | Cancer | T-cells were expanded, sorted, and sequenced against tumor-associated antigens (SIY & TRP2) in the B16 cell line. |
| Dash_2017 | Human & Murine | Infectious Disease | T-cells were sorted and sequenced from either mice or humans for 7 murine and 3 human Class-I specificities. |
| Zhang_2018 | Human | Infectious Disease & Cancer | T-cells were high-throughput screened for a large number of human viral, tumor-associated, and neoantigens via previously published method TetTCR-Seq. Antigens with positivity for ≥ 20 unique TCR's were selected for analysis. |
| 10x_Genomics | Human | Infectious Disease & Cancer | T-cells were high-throughput screened for a large number of human viral and tumor-associated antigens via published 10x Genomics single-cell pipeline. |
| Rudqvist_2017 | Murine | Cancer | Tumor-Infiltrating Lymphocytes (TIL) were collected from tumor-bearing mice who either received no therapy (Control), radiation (RT), α-CTLA4 (9H10), or combination therapy (RT+9H10). 20 mice were sequenced in this study after treatment and tumor growth with 5 mice/treatment cohort. |
| CheckMate-038 | Human | Cancer | Tumor biopsy samples were collected from 43 patients with metastatic melanoma undergoing a clinical trial (CheckMate-038, parts 2-4 of NCT01621490) for either α-PD1 monotherapy (9 patients) or α-PD1+α-CTLA4 combination therapy (34 patients) prior to receiving therapy. TCR β-chain sequencing was conducted on DNA extractions from these biopsies. Patients were selected for analysis who had pre-treatment TCR-Seq, WES sequencing available for HLA genotyping, and clinical response follow-up (as determined by RECIST v1.1) |

**Supplementary Figure 2. Dataset Descriptions.** DeepTCR was piloted on sources of data that covered both human and mouse TCR's including samples taken from infectious disease settings and cancer pathology.

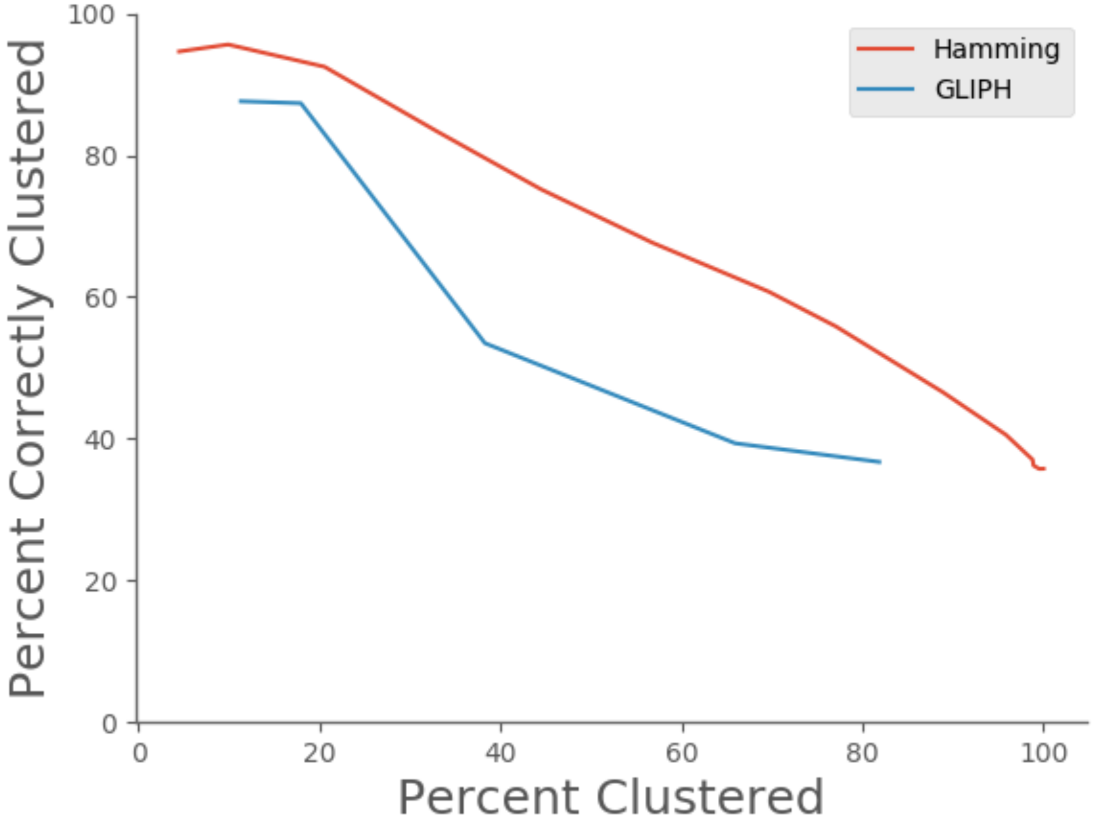


**Supplementary Figure 2. Comparing clustering performance of Hamming Distance v. GLIPH.** In order to assess the performance of a simple Hamming distance vs the state-of-the-art TCR-Seq clustering algorithm, we applied a Hamming distance followed by hierarchical clustering following complete linkage to the *Glanville* dataset of 2066 TCR sequences specific for 7 antigens as well as ran the GLIPH clustering algorithm. We then assessed the clustering accuracy (as per methods used by *Glanville et. al.*). This entailed only including clusters with at least 3 sequences and making cluster assignments based on the majority of members within a cluster. If there was no majority, no cluster assignment was made. For clusters with assignments, sequences within the cluster that shared the cluster assignment labeled were counted as being clustered correctly.

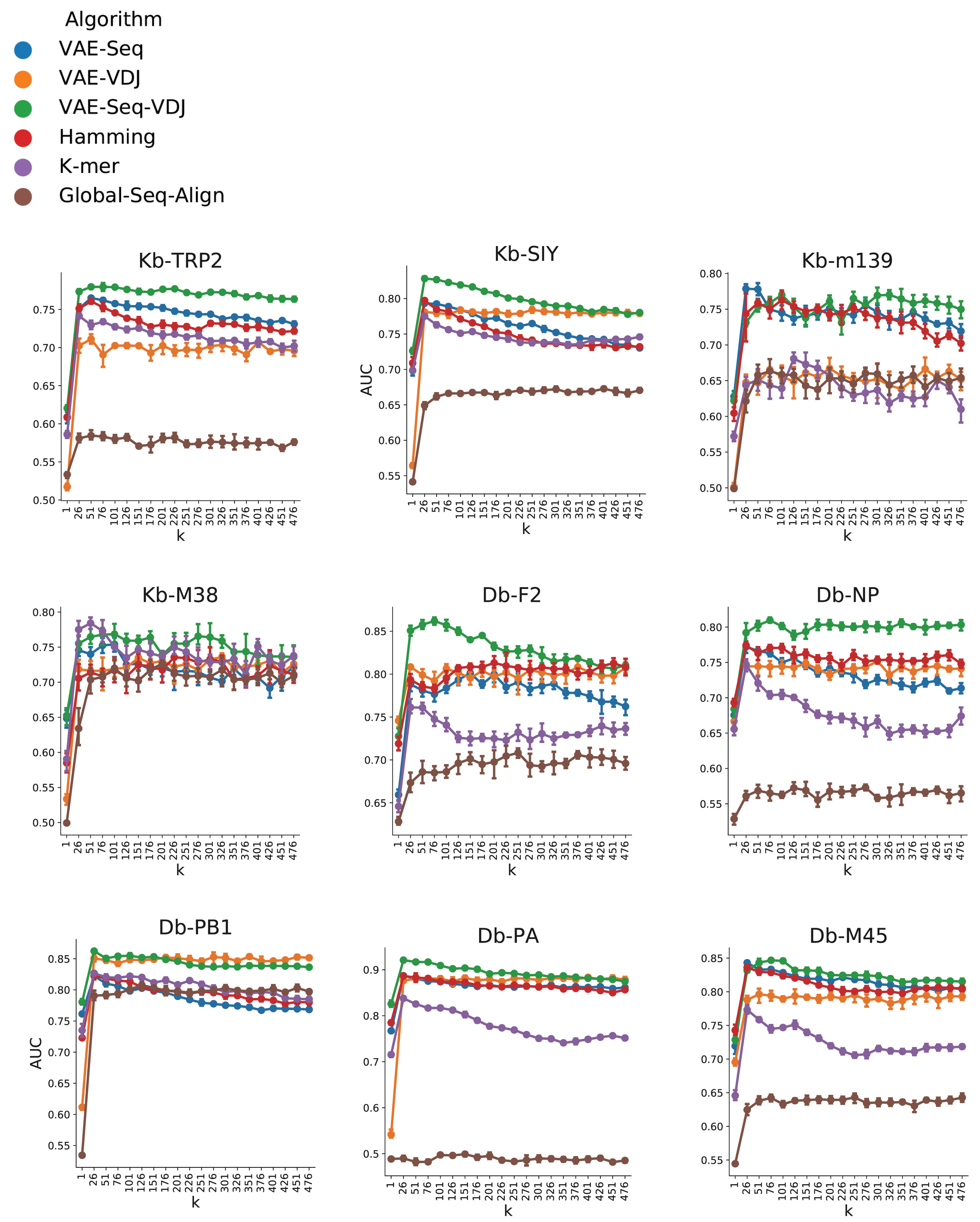


**Supplementary Figure 3. AUC Scores for K-Nearest Neighbors on Murine Antigens.** A K-Nearest Neighbors algorithm was applied using a 5-Fold Cross-Validation across various values for k. AUC performance was assessed across all methods

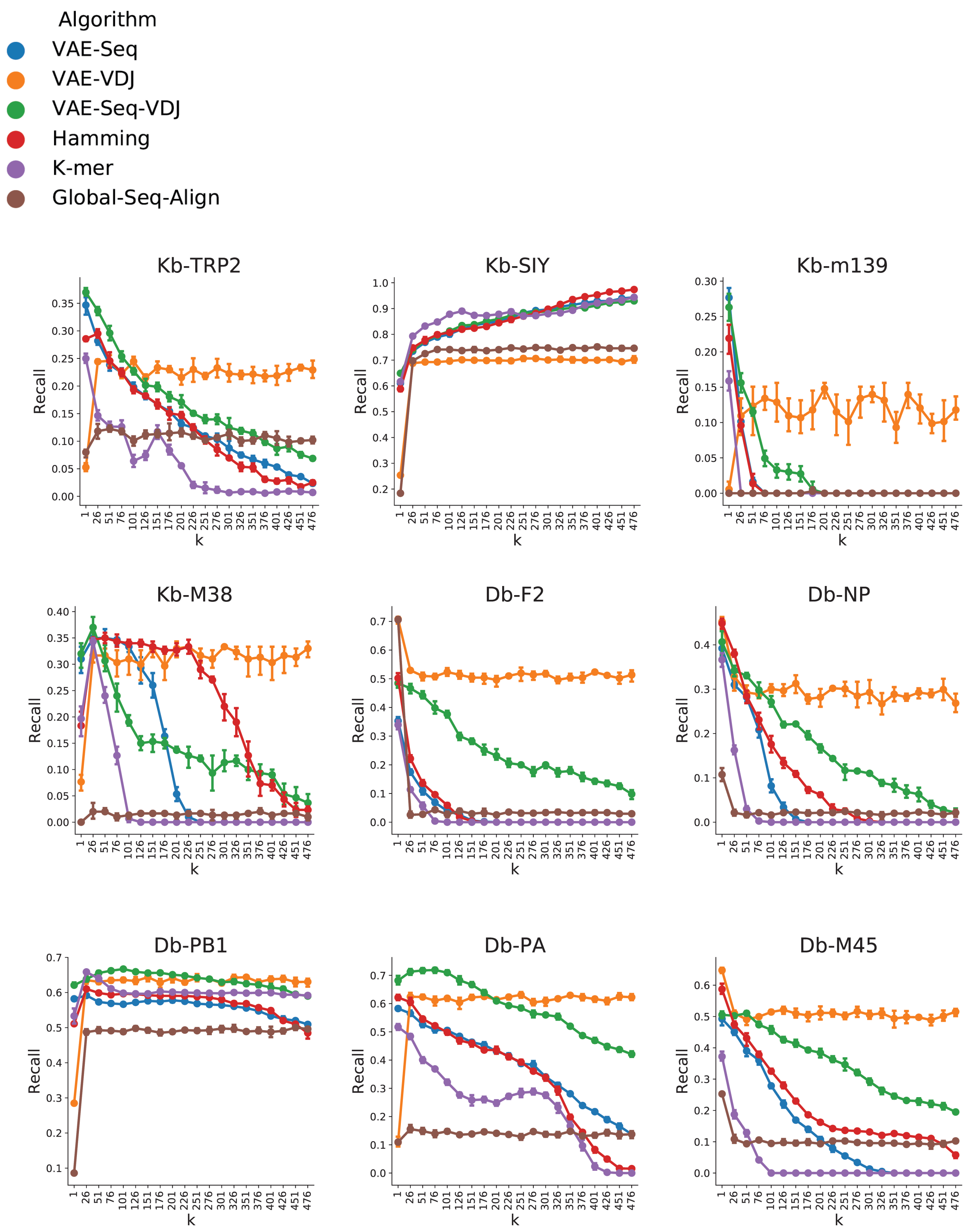


**Supplementary Figure 4. Recall Scores for K-Nearest Neighbors on Murine Antigens.** A K-Nearest Neighbors algorithm was applied using a 5-Fold Cross-Validation across various values for k. Recall performance was assessed across all methods.

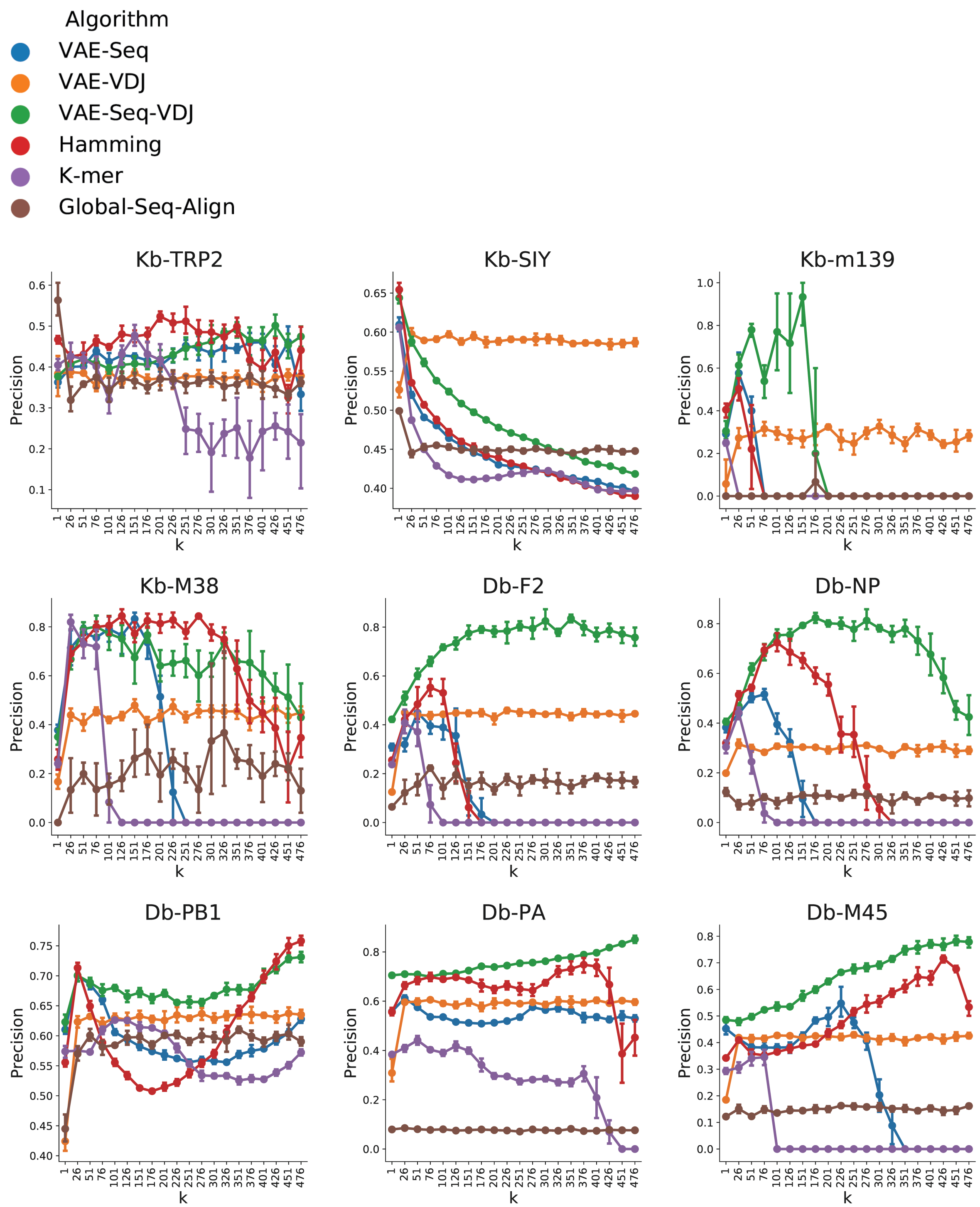


**Supplementary Figure 5. Precision Scores for K-Nearest Neighbors on Murine Antigens.** A K-Nearest Neighbors algorithm was applied using a 5-Fold Cross-Validation across various values for k. Precision performance was assessed across all methods.

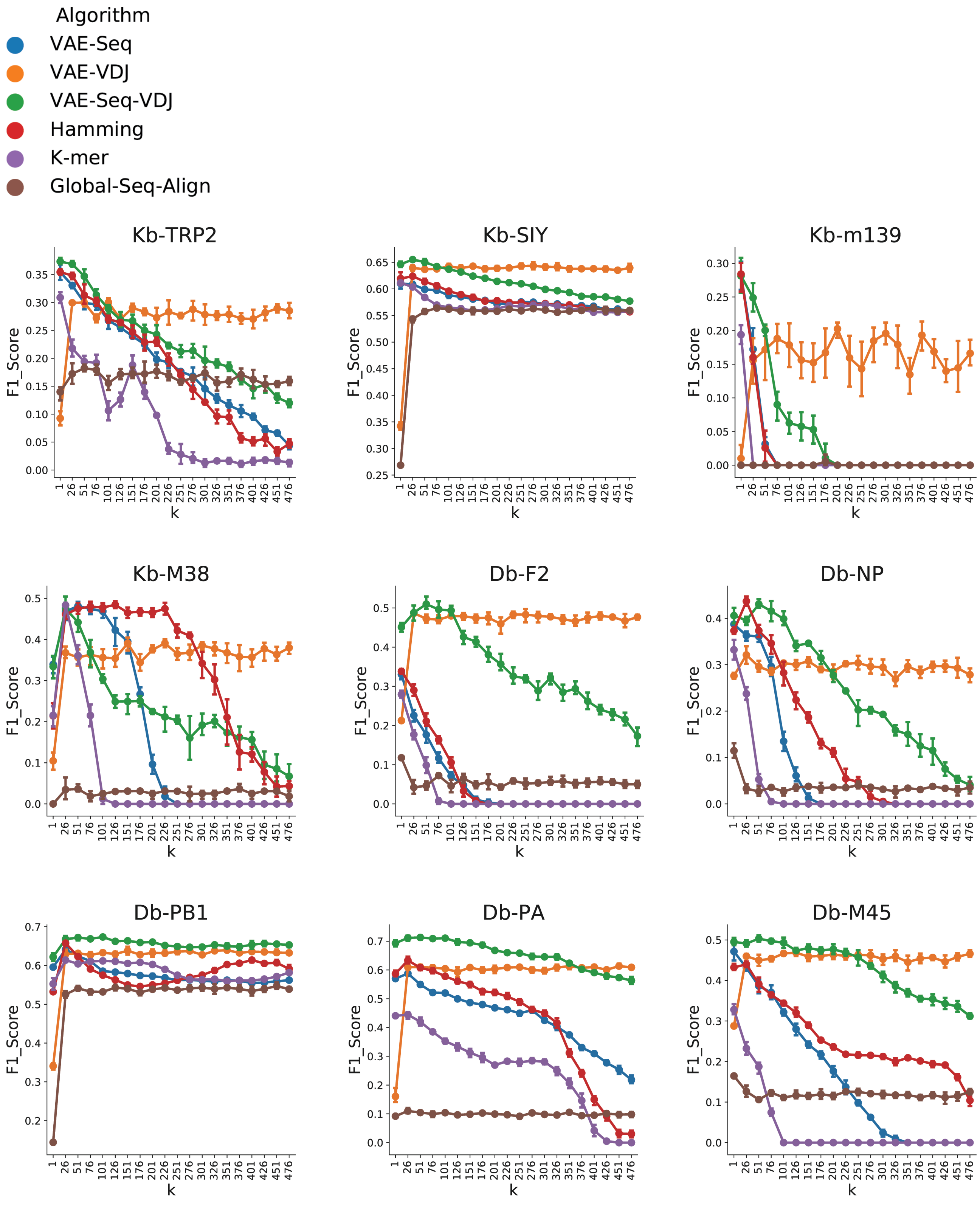


**Supplementary Figure 6. F1 Scores for K-Nearest Neighbors on Murine Antigens.** A K-Nearest Neighbors algorithm was applied using a 5-Fold Cross-Validation across various values for k. F1 Score was assessed across all methods.

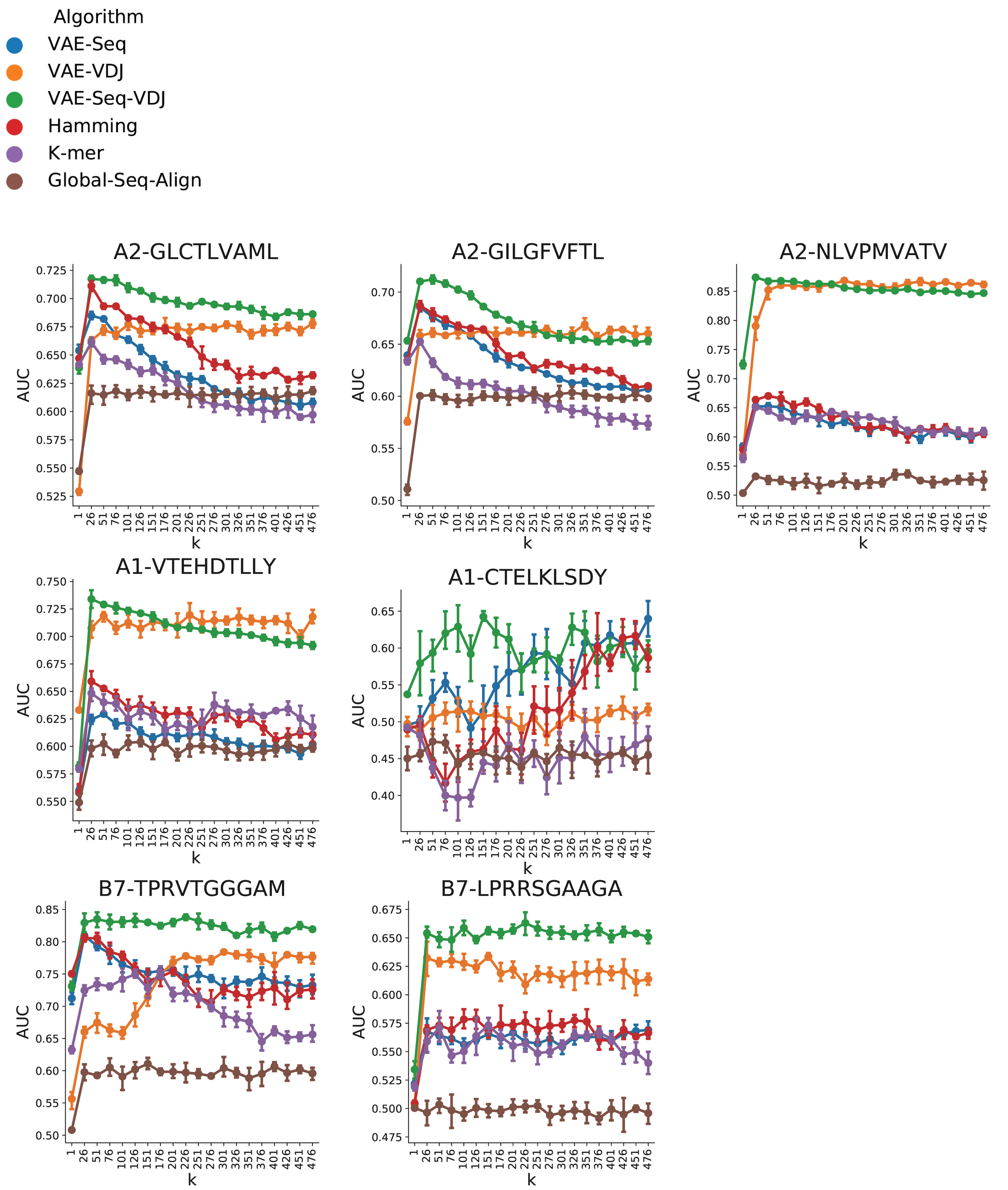


**Supplementary Figure 7. AUC Scores for K-Nearest Neighbors on Human Antigens.** A K-Nearest Neighbors algorithm was applied using a 5-Fold Cross-Validation across various values for k. AUC performance was assessed across all methods.

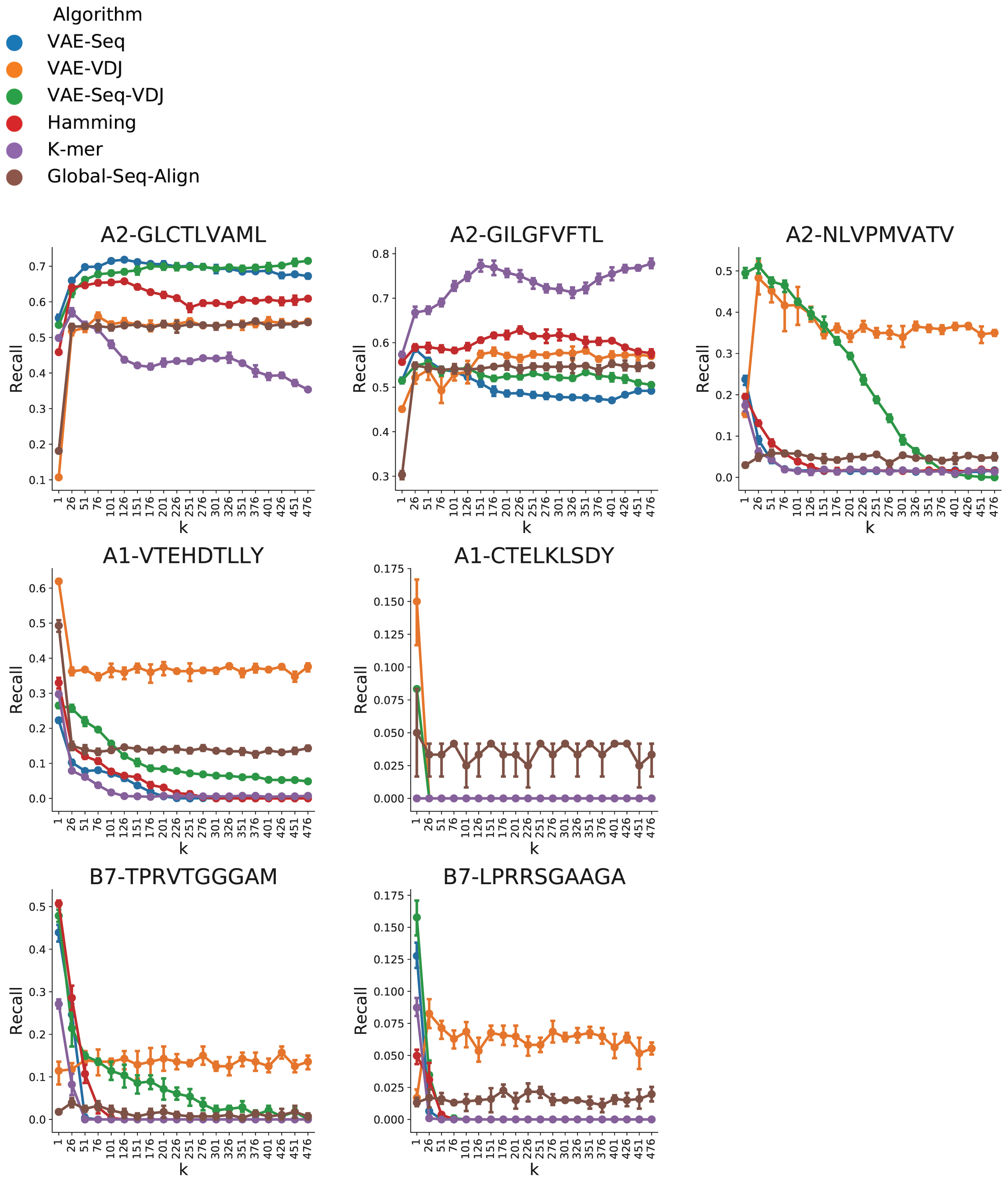


**Supplementary Figure 8. Recall Scores for K-Nearest Neighbors on Human Antigens.** A K-Nearest Neighbors algorithm was applied using a 5-Fold Cross-Validation across various values for k. Recall performance was assessed across all methods.

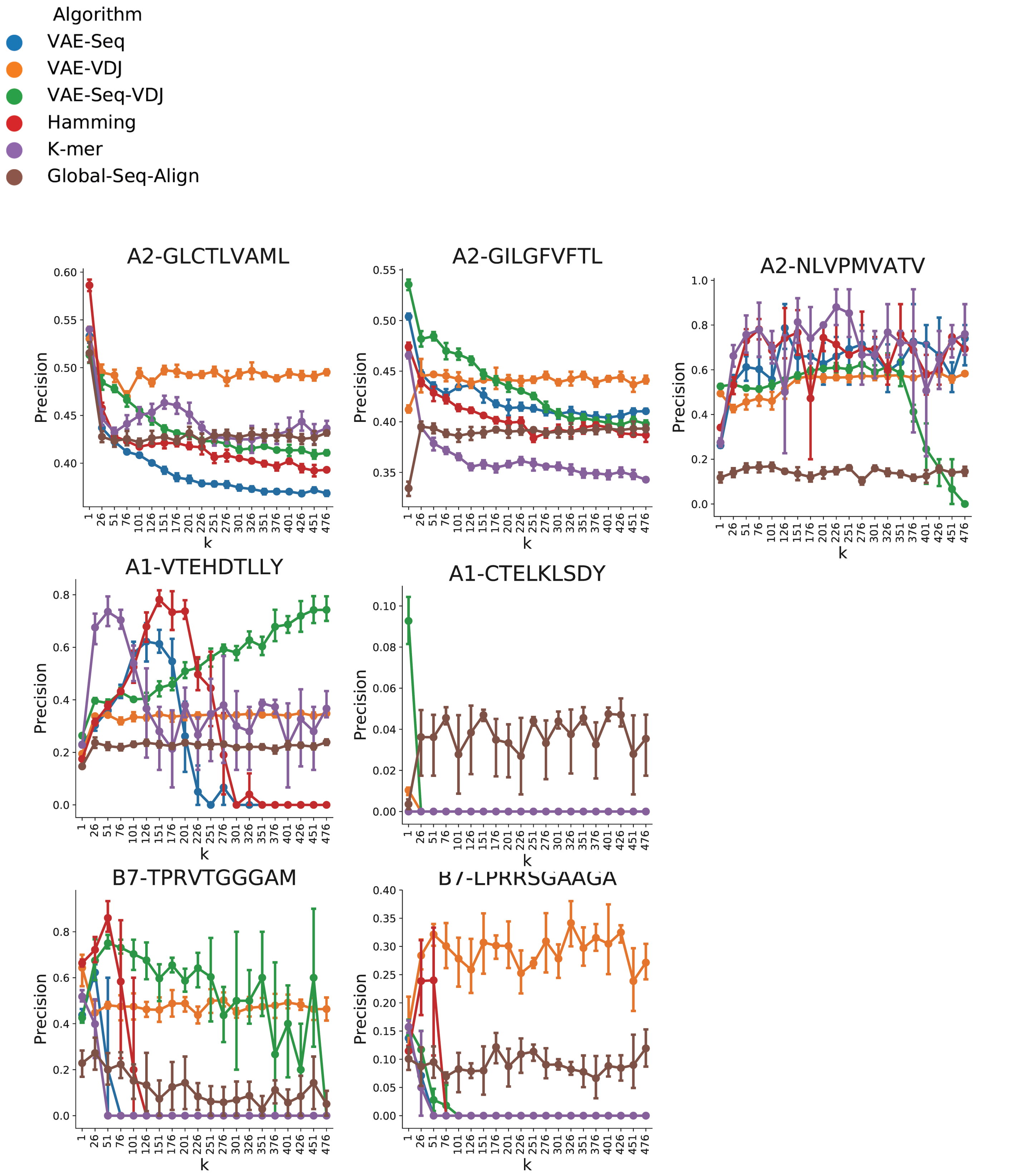


**Supplementary Figure 9. Precision Scores for K-Nearest Neighbors on Human Antigens.** A K-Nearest Neighbors algorithm was applied using a 5-Fold Cross-Validation across various values for k. Precision performance was assessed across all methods.

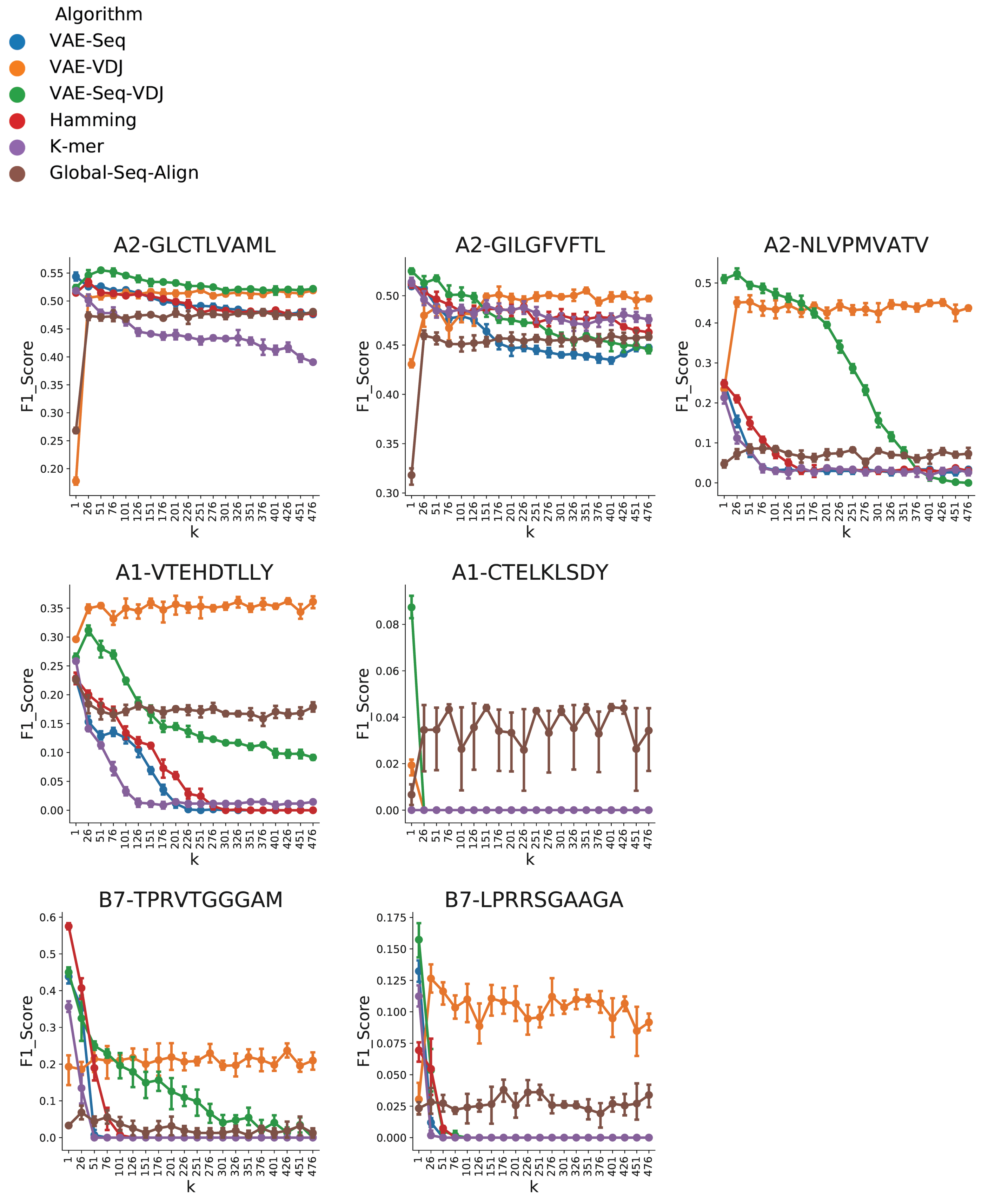


**Supplementary Figure 10. F1 Scores for K-Nearest Neighbors on Human Antigens.** A K-Nearest Neighbors algorithm was applied using a 5-Fold Cross-Validation across various values for k. F1 Score was assessed across all methods.

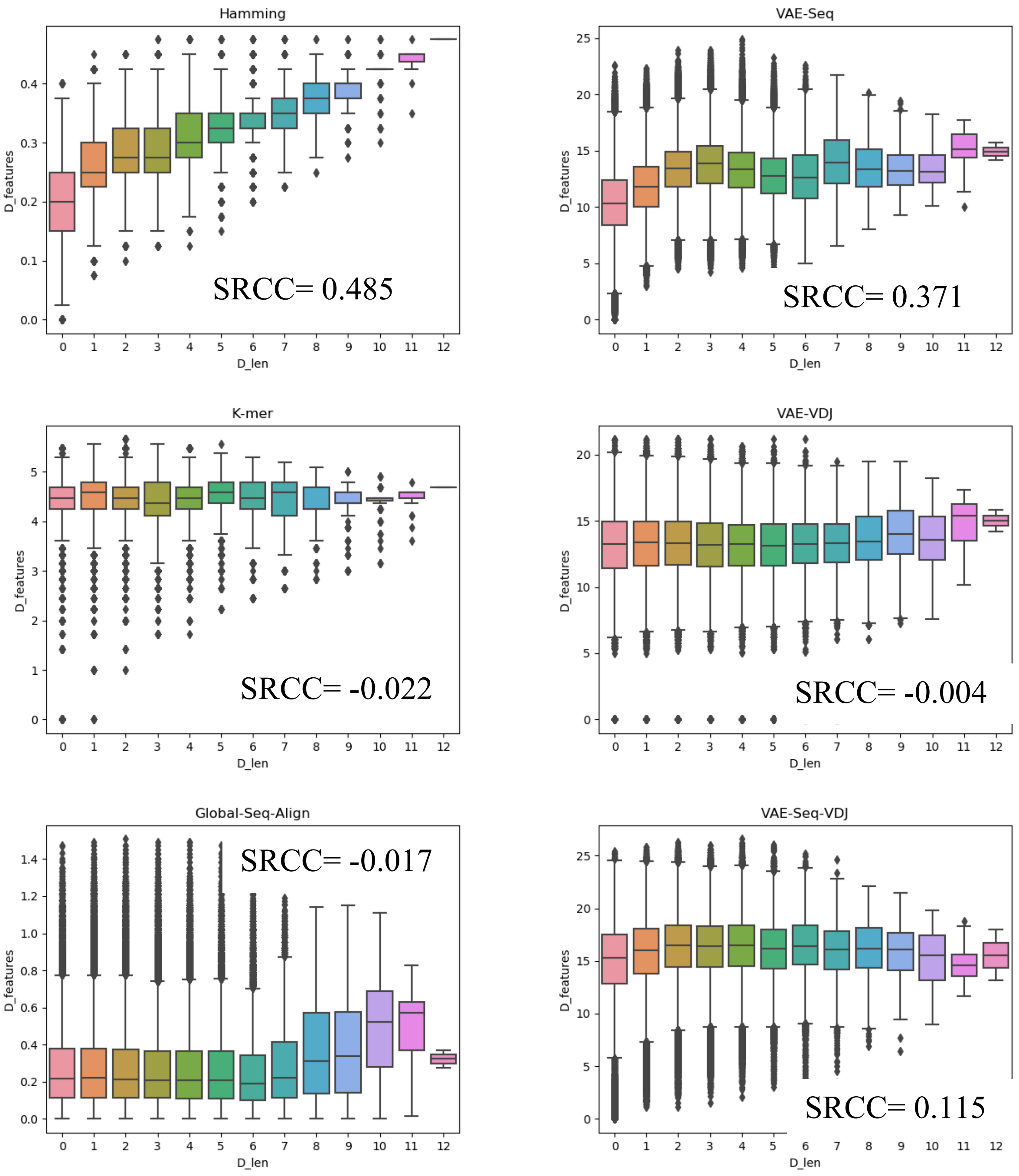


**Supplementary Figure 11. Assessing correlation between various distance metrics and length of the sequence.** In order to assess the extent by which these various methods used to quantify the distance between sequences was driven by sequence length, we determined the Spearman's Rank Correlation Coefficient (SRCC) between the distances of a given pair of sequences and how different in length they were. Boxplots are shown to compare distance by sequence length (x-axis) by distance of various methods (y-axis).

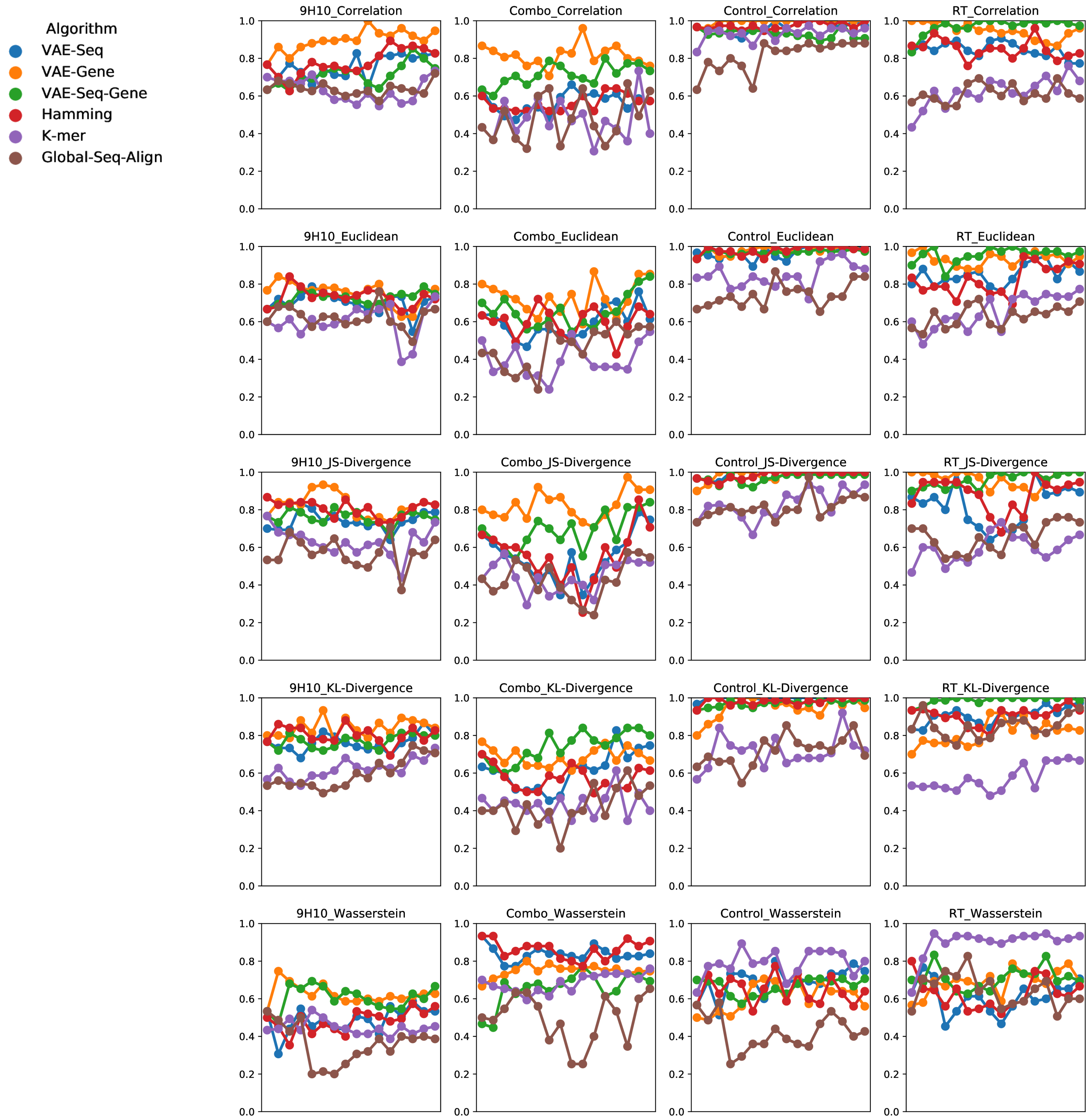


**Supplementary Figure 12. AUC Scores for K-Nearest Neighbors on samples taken from murine tumor-infiltrating lymphocytes (TIL).** A K-Nearest Neighbors algorithm was applied using a 5-fold K-fold Cross-Validation using k-values from 1-16 for classifying a given repertoire given the distribution of its repertoire across the PhenoGraph clustering algorithm solution. Various distance metrics were used to compare the distance between these empirical distributions. Scores were assessed across all sequence distance methods.

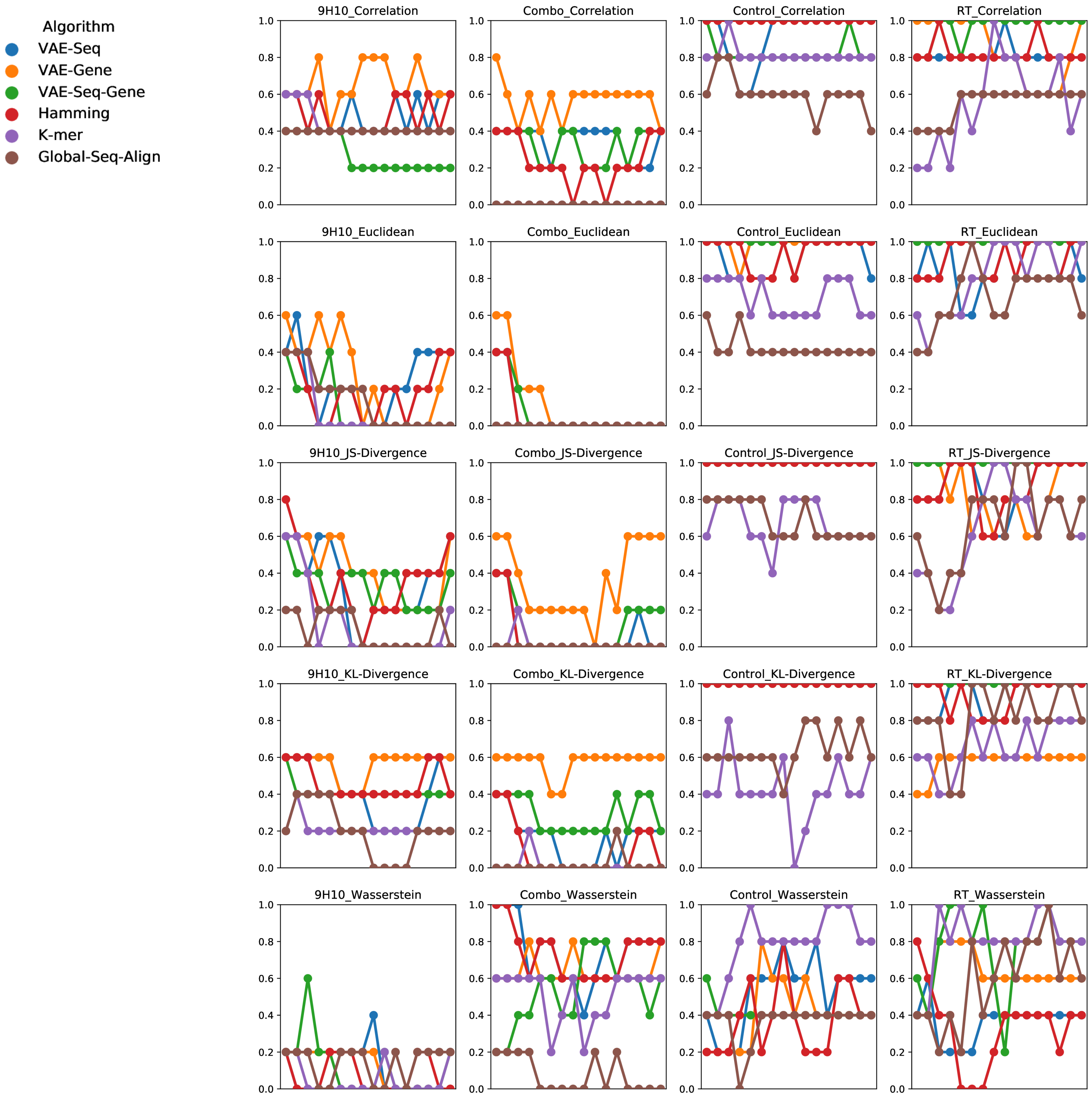


**Supplementary Figure 13. Recall Scores for K-Nearest Neighbors on samples taken from murine tumor-infiltrating lymphocytes (TIL).** A K-Nearest Neighbors algorithm was applied using a 5-fold K-fold Cross-Validation using k-values from 1-16 for classifying a given repertoire given the distribution of its repertoire across the PhenoGraph clustering algorithm solution. Various distance metrics were used to compare the distance between these empirical distributions. Scores were assessed across all sequence distance methods.

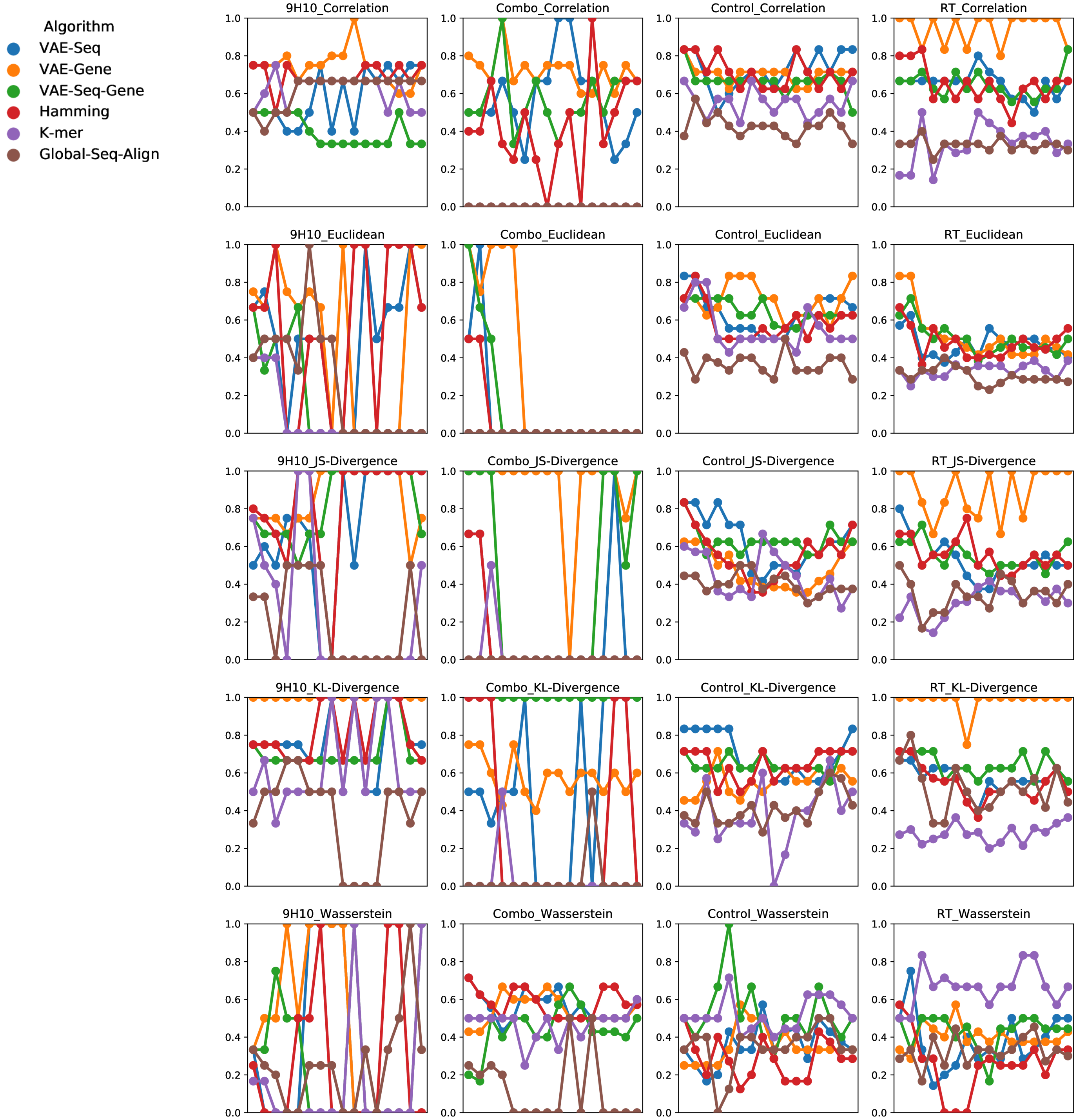


**Supplementary Figure 14. Precision Scores for K-Nearest Neighbors on samples taken from murine tumor-infiltrating lymphocytes (TIL).** A K-Nearest Neighbors algorithm was applied using a 5-fold K-fold Cross-Validation using k-values from 1-16 for classifying a given repertoire given the distribution of its repertoire across the PhenoGraph clustering algorithm solution. Various distance metrics were used to compare the distance between these empirical distributions. Scores were assessed across all sequence distance methods.

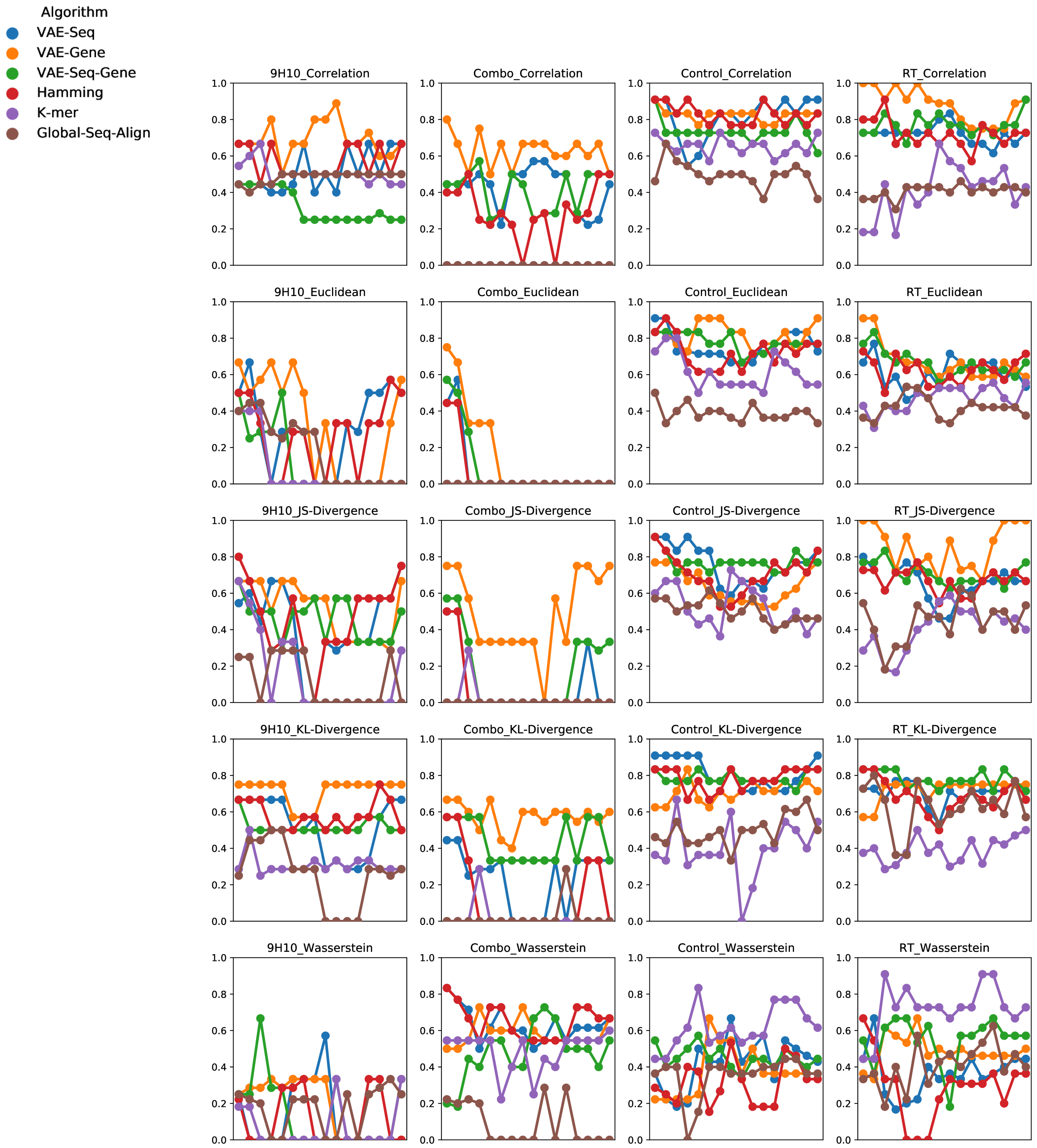


**Supplementary Figure 15. F1 Scores for K-Nearest Neighbors on samples taken from murine tumor-infiltrating lymphocytes (TIL).** A K-Nearest Neighbors algorithm was applied using a 5-fold K-fold Cross-Validation using k-values from 1-16 for classifying a given repertoire given the distribution of its repertoire across the PhenoGraph clustering algorithm solution. Various distance metrics were used to compare the distance between these empirical distributions. Scores were assessed across all sequence distance methods.

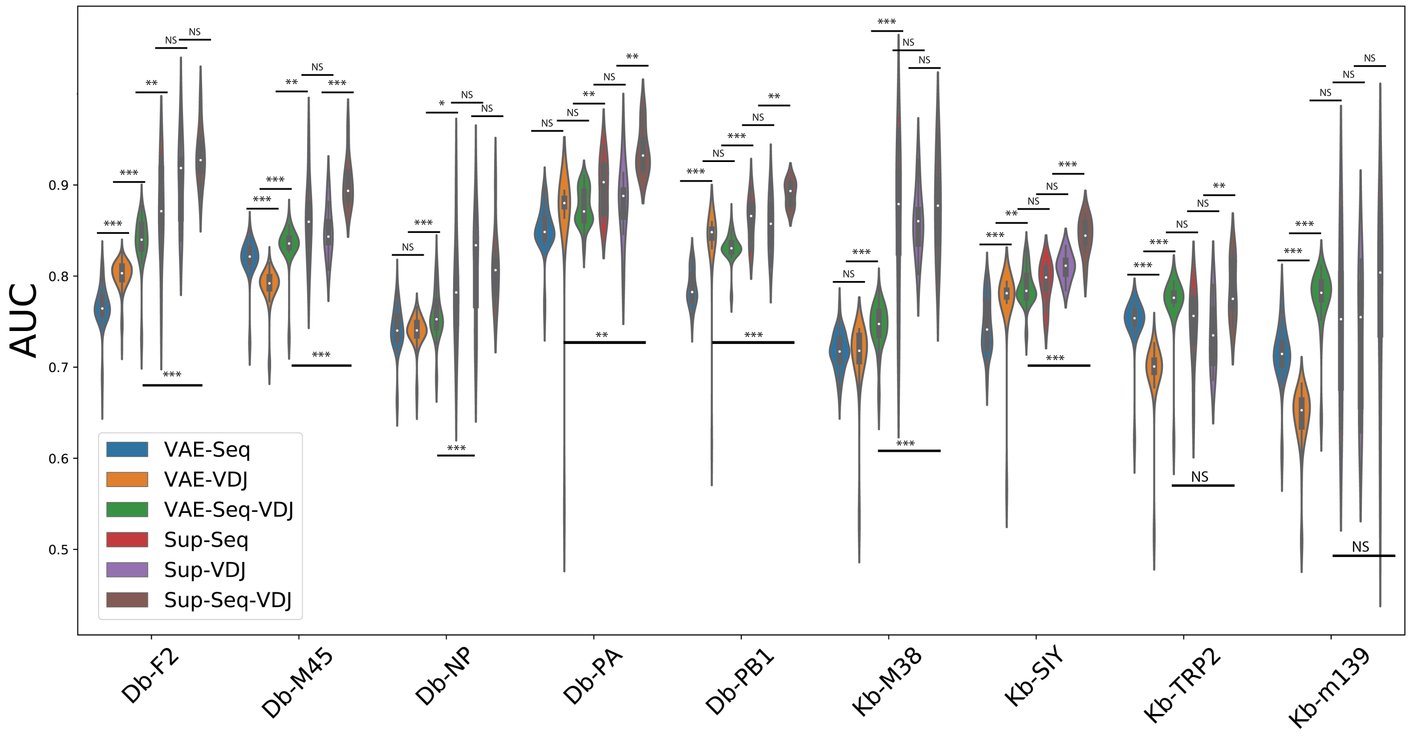


**Supplementary Figure 16. Benchmarking Classification Performance between unsupervised and supervised deep learning methods.** AUC for KNN sequence classifiers for unsupervised representations via VAE vs supervised sequence classifiers. (Two-sample independent t-test, * : p < 0.05, ** : p < 0.01, *** : p < 0.001)

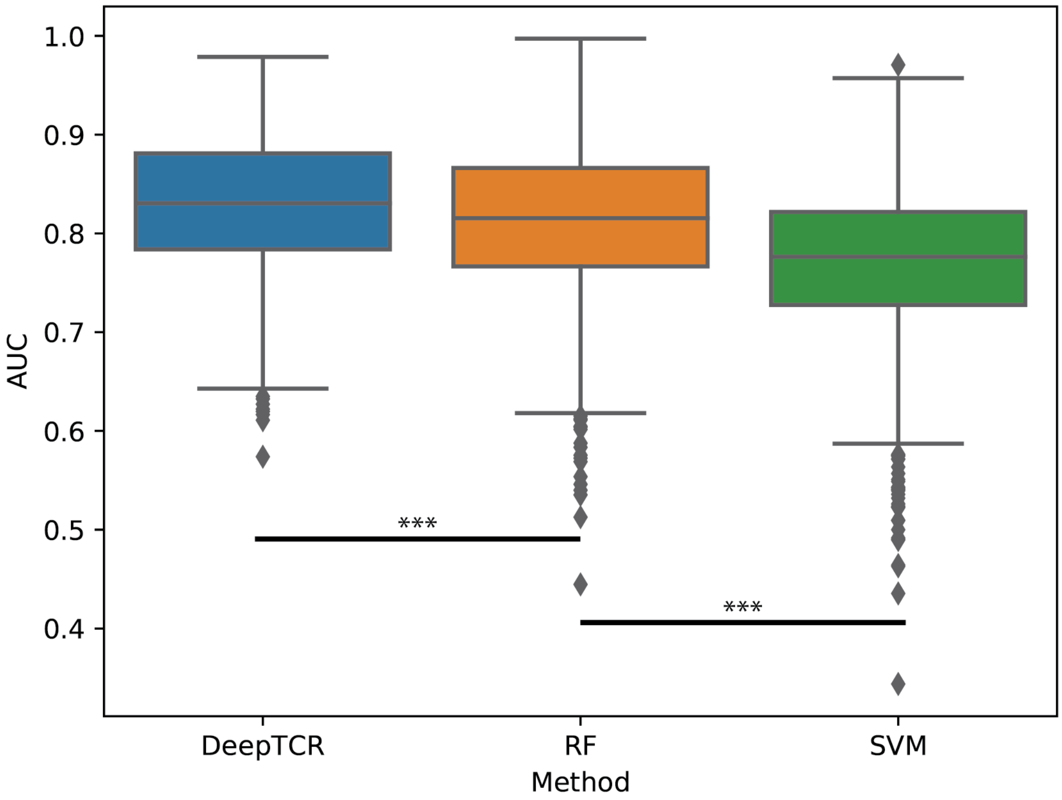


**Supplementary Figure 17. Benchmarking various machine learning methods to classify TCR sequences by their antigen-specificity.** In order to compare the performance of various machine learning approaches to classify TCR sequences by their antigen-specificity, we used the TCR sequences for the 9 murine antigens collected to benchmark DeepTCR's deep learning classifier vs a classical Support Vector Machine (SVM) and Random Forrest (RF) classifier. Only the beta cdr3 sequence was provided to all 3 machine learning algorithms to test the ability of the classifiers to predict antigen-specificity from cdr3 motifs. 100 iterations of a 5-fold cross-validation were completed and AUC's were measured for each of the various methods to assess classification performance. Average AUC values for DeepTCR's deep learning sequence classifier, the Random Forrest, and SVM were 0.830, 0.810, and 0.769 respectively. (Two-sample independent t-test, * : $p < 0.05$, ** : $p < 0.01$, *** : $p < 0.001$)

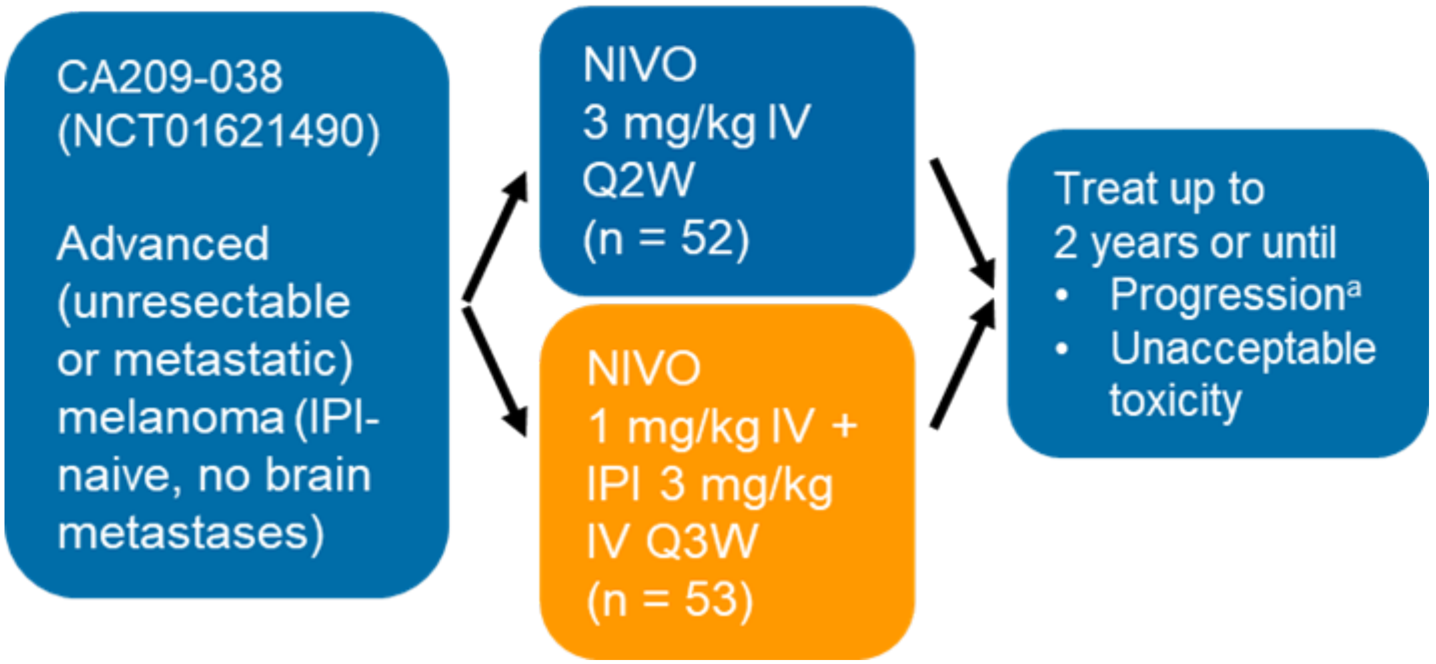


**Supplementary Figure 18. CheckMate-038 Clinical Trial Schema.** Patients from parts 2-4 of CheckMate-038 with metastatic melanoma were randomized into either α-PD1 monotherapy (9 patients where data was available) or α-PD1 + α-CTLA4 combination therapy (34 patients where data was available). Patients were followed up 6 months after beginning therapy via assessing radiographic response via RECIST v1.1. TCR-Seq was performed on pre-therapy tumor biopsies. HLA genotyping was determined from whole exome sequencing. Full details of trial design can be found at ClinicalTrials.gov (NCT01621490).

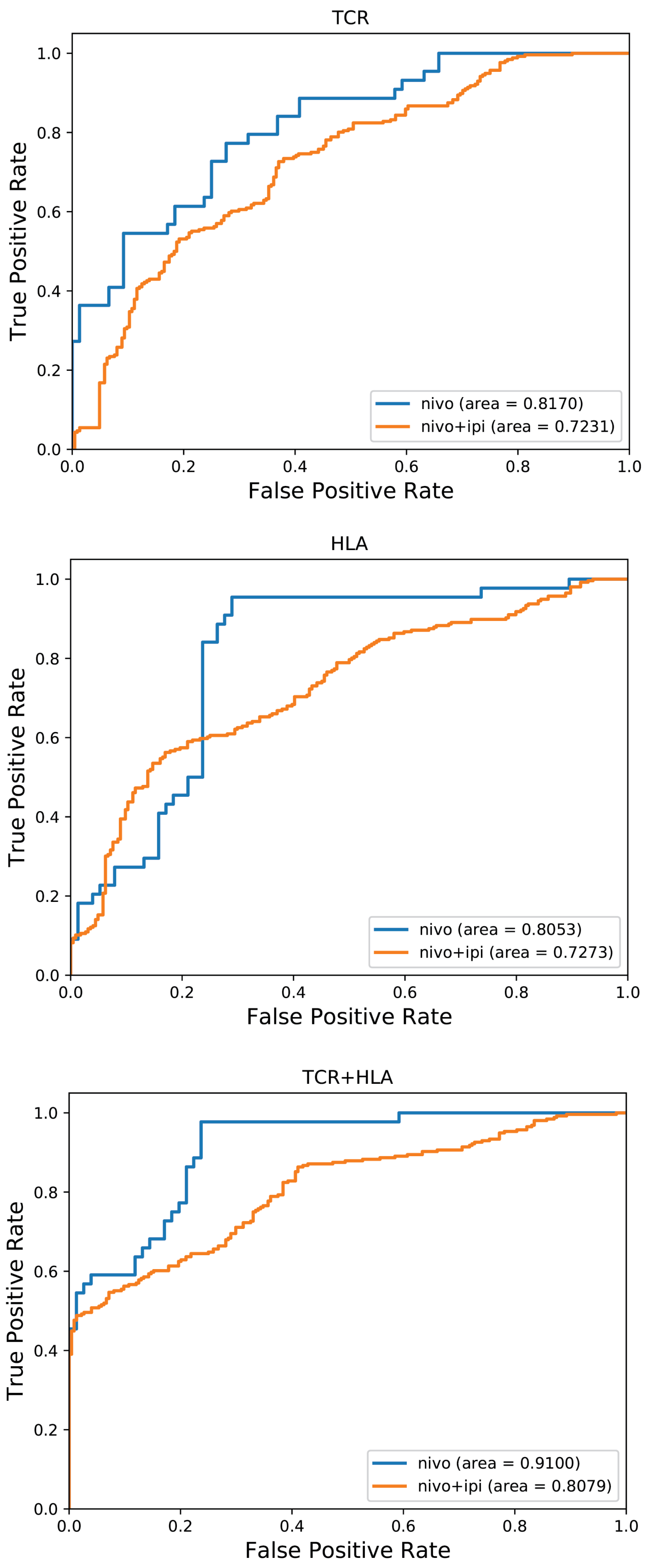


**Supplementary Figure 19. DeepTCR Performance on CheckMate-038 data stratified by treatment cohort.** Data from Fig. 3d was stratified by treatment cohort and ROC performance was assessed for the described DeepTCR models (TCR, HLA, TCR+HLA).

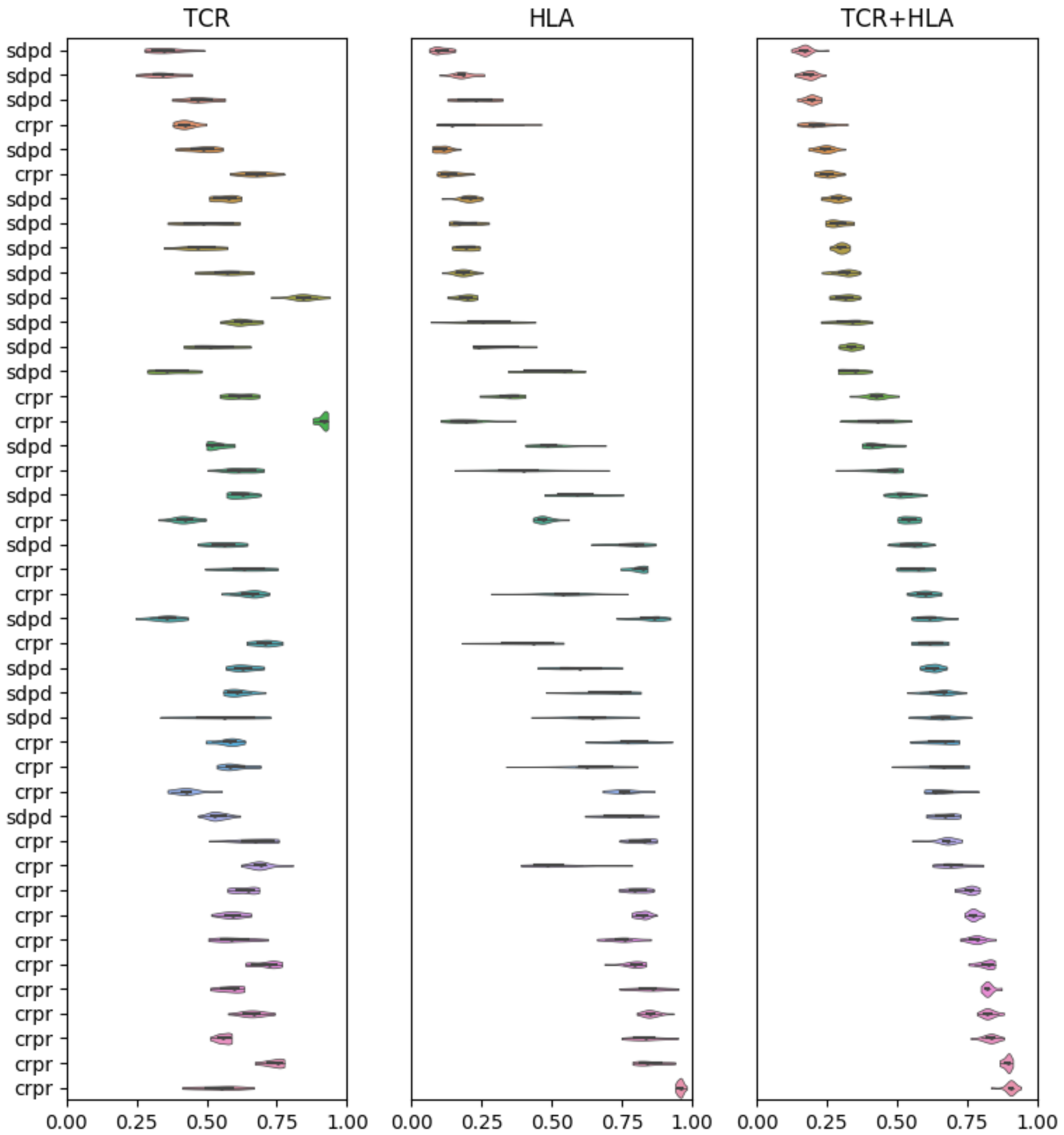


**Supplementary Figure 20. Distribution of CheckMate-038 Sample Predictions from Various Models.** Predictions from Monte-Carlo simulations were collected across all trained models (TCR, HLA, TCR+HLA) and the distribution of predictions for each sample was plotted to observe not only the mean prediction but the distribution of predictions per sample to assess the certainty of the model to predict any individual sample. It was noted that the TCR+HLA model generates predictions that have a smaller confidence interval and thus, are more certain than either the TCR or HLA models (Wilcox signed-rank test:: HLA vs TCR+HLA: p-val = 4.73e-6, TCR vs TCR+HLA: p-val = 5.70e-3)

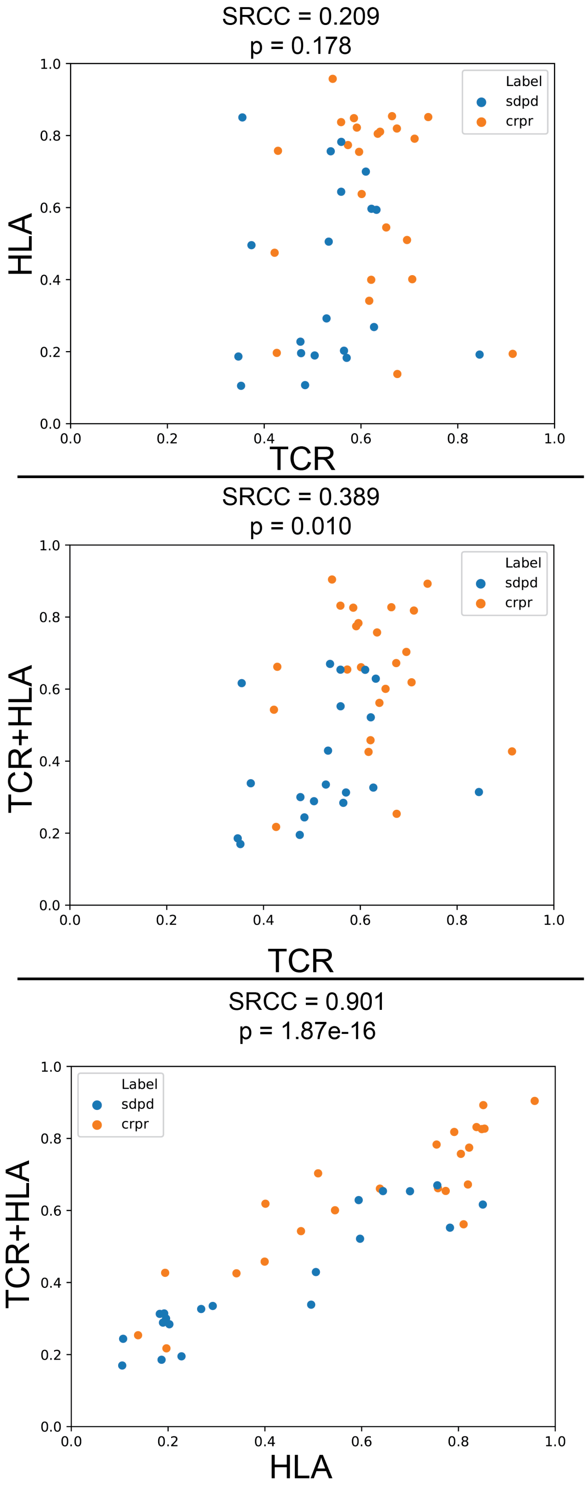


**Supplementary Figure 21. Sample Predictions for CheckMate-038 trial patients from different repertoire classification models.** To assess the shared information between data provided by HLA of the patients vs their TCR repertoire, we took the average predicted value for a given sample over 100 Monte-Carlo simulations from the 3 repertoire models (TCR information alone, HLA information alone, and TCR+HLA information combined), and computed the Spearman's rank correlation coefficient (SRCC) of these prediction values across all pairwise combinations of models. These results suggest that HLA conditions the information that is learned from the TCR repertoire when used in conjunction to train the model as there is little correlation between HLA and TCR models alone but high correlation between HLA and TCR+HLA models.

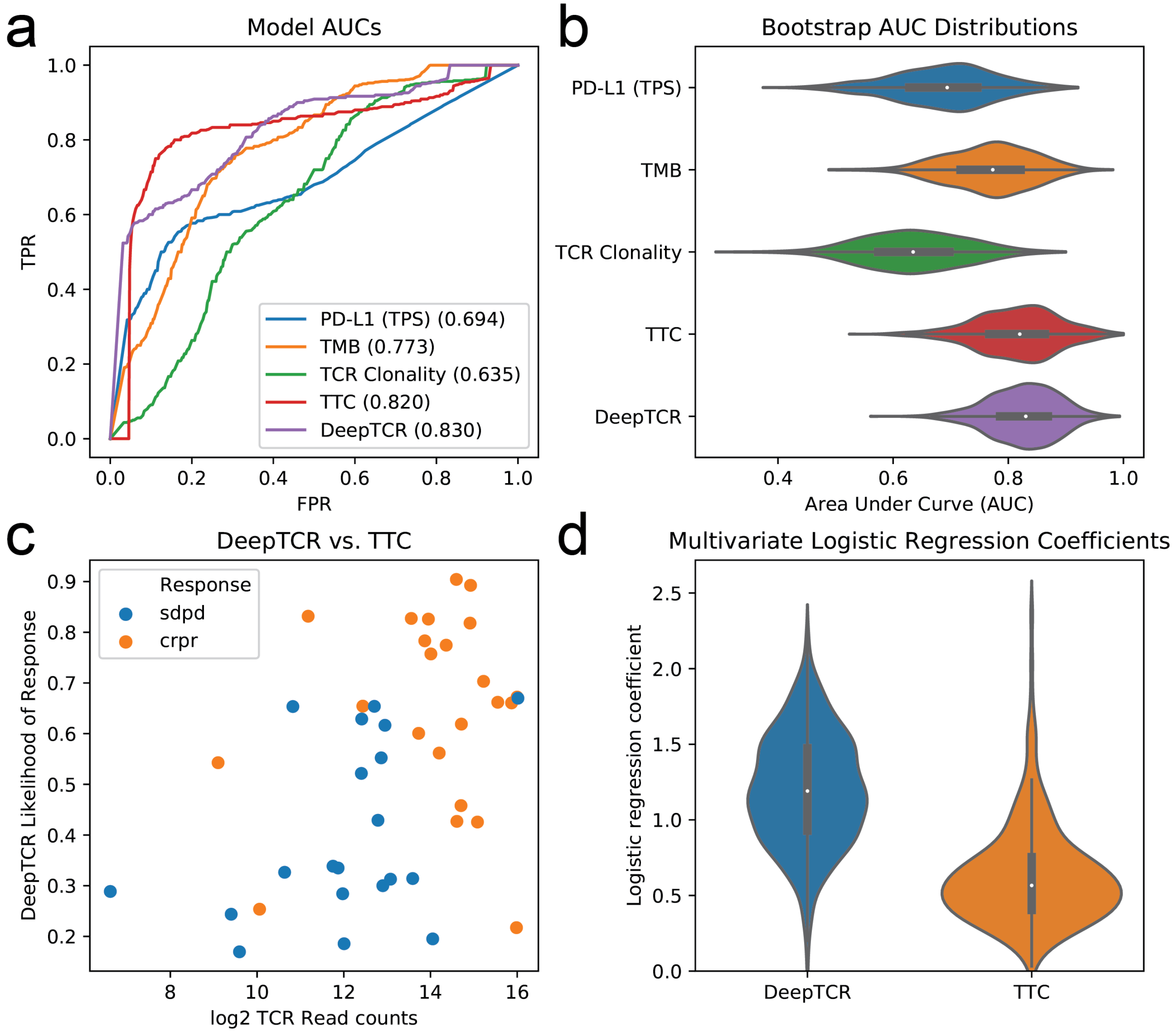


**Supplementary Figure 22. Comparison of DeepTCR against other biomarkers in CheckMate-038. (a)** In order to compare predictive performance of DeepTCR against other conventional biomarkers often used to stratify response in immunotherapy-treated patients, we took the average sample prediction from the Monte-Carlo simulations (Fig. 3c) and compared area under the curve (AUC) from the receiver operating characteristic curves of DeepTCR against other TCR sequence derived metrics, TCR Clonality and total T-cell count (TTC), along with immunohistochemistry (IHC) of PD-L1, tumor positive score (TPS), and exome based total mutational burden (TMB). **(b)** Bootstrap estimates were used to characterize the distribution of AUC for each biomarker as well as compare the difference between pairs of biomarkers. **(c)** DeepTCR likelihood of response predictions were compare against TTC (spearman rank correlation coefficient = 0.44). **(d)** When DeepTCR and TTC are used in multivariate logistic regression modeling, they remain as two independent predictors as evidenced by the fact that the 95% bootstrapped confidence intervals of the model coefficients do not cross 0 (95% CI; DeepTCR = [0.486, 1.817], TTC = [0.185, 1.353]).

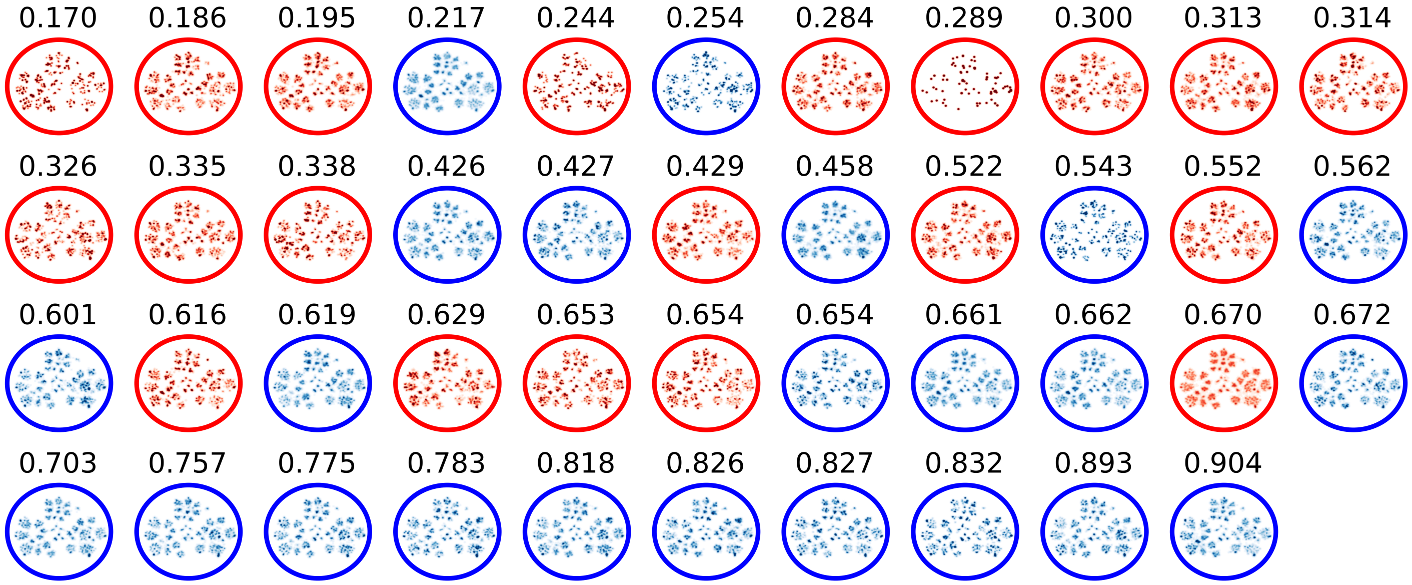


**Supplementary Figure 23. UMAP representation of unsupervised VAE featurization for all sequence data in CheckMate-038.** All sequence data from the CheckMate-038 clinical trial was featurized (TCR+HLA information) via an unsupervised VAE and UMAP representations were created on a per sample basis to show the unfiltered distribution of data.
